# Molecular epidemiology of Cercospora leaf spot on resistant and susceptible sugar beet hybrids

**DOI:** 10.1101/2023.10.30.564591

**Authors:** Chen Chen, Harald Keunecke, Enzo Neu, Friedrich J. Kopisch-Obuch, Bruce A. McDonald, Jessica Stapley

## Abstract

Cercospora leaf spot (CLS), caused by *Cercospora beticola*, is a major foliar disease impacting sugar beet production worldwide. The development of new resistant sugar beet hybrids is a powerful tool to better manage the disease, but it is unclear how these hybrids affect CLS epidemiology. We used a molecular epidemiology approach to study natural epidemics of CLS affecting two susceptible and two resistant sugar beet hybrids at two field sites.

Infected plants were geotagged on a weekly basis. Isolations of *C. beticola* were made from infected leaves and genotyped using six simple sequence repeat loci to identify clones. We determined that CLS epidemics had a later onset in plots planted to resistant hybrids, but once the pathogen established an infection, there was little difference between resistant and susceptible hybrids in the probability of localized spread and dispersal. We found that different clones often infected the same leaf and that clusters of infected plants were often colonized by a mixture of clones. There was little overall difference in genetic diversity between resistant and susceptible hybrids, however genotypic evenness differed at one site. In this site we found one genotype restricted to the resistant cultivars at a high frequency. At the end of the epidemic infections were not randomly distributed across the fields and we found that a single clone could spread over a distance of 100 m during a growing season.

## Introduction

Sugar beet (*Beta vulgaris* L.) contributes approximately one-fifth of global sucrose production annually (Rangel et al., 2020). The most destructive disease of sugar beet is the foliar disease Cercospora leaf spot (CLS) caused by the fungus *Cercospora beticola*. CLS affects the production of sugar beet in most sugar beet growing regions worldwide (Holtschulte, 2000). Under high disease pressure, the yield loss can exceed 40% (Shane and Teng, 1992). The main method of control is multiple applications of fungicides, however this has resulted in the emergence of widespread fungicide resistance in *C. beticola* populations (Bolton et al., 2013; Kayamori et al., 2021). Resistant cultivars are seen as an important means of control in addition to fungicides, with CLS-resistant cultivars providing protection under low to moderate disease pressure (Francis and Luterbacher, 2003; Vogel et al., 2018; Weiland and Koch, 2004; Wolf and Verreet, 2005). A new resistance gene named *BvCR4* has recently been identified and bred into sugar beets (Törjék et al., 2020). This new gene provides a high level of CLS resistance and could be a useful method of control.

As with the evolution of fungicide resistance, pathogens can evolve to overcome host resistance through selection for virulent strains. How quickly pathogens evolve virulence to resistant cultivars is strongly influenced by the genetic architecture of the resistance – single major resistance (R) genes with large effect are usually less durable compared to quantitative resistance encoded by many genes (Cowger and Brown, 2019; Gou et al., 2023). The evolution of virulence is also influenced by the amount of genetic diversity in the selected pathogen population, the deployment strategies used for the resistance genes, and the distribution of virulent pathogen genotypes across a heterogenous host population (Milgroom and Peever, 2003; Rimbaud et al., 2018; Stam and McDonald, 2018).

The interaction between pathogen genotypes and host genotypes shapes both the development of epidemics at the field scale and the evolutionary landscape of the entire pathosystem at larger spatial and temporal scales. High genetic diversity within a pathogen population can promote the evolution of virulence (McDonald and Linde, 2002; Sacristán et al., 2021), but the type and degree of host resistance can influence the distribution of genetic diversity in the corresponding pathogen population. When a single new resistant cultivar carrying a major R gene is first deployed in the field, the corresponding pathogen population experiences strong selection and a bottleneck, because only genotypes that carry the compatible virulence factors can infect the resistant cultivar. This selective bottleneck can result in a reduction in genetic and genotypic diversity in the pathogen population. How quickly the pathogen population and the corresponding genetic diversity can recover following this selection bottleneck depends upon the life history and evolutionary potential of the pathogen - in particular the frequency of sexual reproduction and the efficiency of dispersal of virulent genotypes (Barrett et al., 2008; McDonald and Linde, 2002). Though a sexual stage (teleomorph) has not yet been described for *C. beticola*, population genetic evidence is consistent with regular cycles of sexual recombination. This evidence includes the presence of both mating types at the *MAT* locus, often in a 1:1 ratio (Bolton et al., 2012; Groenewald et al., 2008), high genotypic diversity and low clonality assessed with neutral genetic markers, and random associations among neutral loci (Groenewald et al., 2008; Vaghefi et al., 2017b). The rapid evolution of fungicide resistance in CLS indicates that *C. beticola* has a high evolutionary potential, suggesting that *C. beticola* populations could also adapt quickly to deployment of resistant host genotypes.

The epidemiology of CLS has been well studied (Imbusch et al., 2021; Rossi et al., 2000; Tedford et al., 2018; Wolf and Verreet, 2005). *C. beticola* is polycyclic and leaf lesions can sporulate after 3 days under ideal conditions (Rossi et al., 2000). The primary site of infection is leaves, CLS infections did not establish on roots and stems inoculated with spores (Khan et al., 2008). The primary inoculum can come from multiple sources, including overwintering structures called pseudostromata, which can persist on plant debris for 22 months (Khan et al., 2008), conidia produced on alternative hosts growing in or nearby new fields, and infested seeds (Spanner et al., 2022). Previous work in the USA found that even when there is no crop rotation, genotypes were not resampled over time, suggesting that the dominant source of inoculum for an epidemic is external to the field of interest (Knight et al., 2018). Conidia can be dispersed to adjacent plants by rainsplash, wind and insects (Khan et al., 2008; Vereijssen et al., 2007; Weiland and Koch, 2004), with the longest recorded dispersal distance exceeding 45 m (Imbusch et al., 2021). How dispersal is influenced by host genotype has not been experimentally tested, but other traits have been considered. On more resistant hosts, spores were smaller, germinated more slowly, and spore yield was reduced, but latency did not vary with host resistance (Rossi et al., 2000). This finding illustrates that resistant hosts may influence CLS epidemiology, which may in turn affect the emergence and distribution of virulent genotypes.

Previous work established that *C. beticola* has high genetic diversity and a high evolutionary potential, and that host resistance may influence aspects of CLS epidemiology, but few studies have investigated the evolution of virulence in *C. beticola* populations or used neutral genetic markers to understand how CLS epidemics develop in fields during a growing season. Molecular epidemiology bridges the gaps between disease epidemiology and pathogen population genetics. By including genetic markers into epidemiology studies, new research questions can be addressed around the origins of epidemics, genotype movements, dispersal pathways and the spatio-temporal aspects of disease progression (Archie et al., 2009; Desbiez et al., 2002; Xhaard et al., 2012).

Here we analyzed the molecular epidemiology of natural CLS epidemics occurring in resistant and susceptible hybrids of sugar beet to better understand the evolutionary potential of this pathogen on newly developed hybrids. The main objects were 1) to determine the time of onset of CLS epidemics on different hybrids; 2) to measure the local dispersal pattern for both primary and secondary inoculum of CLS; 3) to assess the genetic diversity of *C. beticola* populations during epidemics across different spatial and temporal scales, including: on the same leaves; on neighboring plants; and across hybrids within and between fields; 4) to determine the degree of clonality on different host populations and identify the most likely sources of primary and secondary inoculum in natural field infections.

## Methods

### Field trials in Switzerland

Two trial sites located in Switzerland (Rudolfingen, Trüllikon 47°38’18.6"N, 8°40’46.1"E and Hendschiken, Othmarsingen 47°23’40.4"N, 8°13’41.1"E) were planted with four sugar beet hybrids in March 2020. The fields had not been planted with sugar beet in the previous three years. The four hybrids differed in their resistance to CLS, including two highly resistant hybrids with the *BvCR4* gene – Hybrid_R2 and Hybrid_R3, and two susceptible hybrids lacking *BvCR4*– Hybrid_S2 and Hybrid_S3 (Table 1). In each field we selected an experimental area ∼100m long, within the field, that contained plots (12-18 rows) of each of the four hybrids (Figure S1). In Hendschiken, each hybrid was planted in blocks of 12 rows for a total area of 1950 m^2^. In Rudolfingen, each hybrid was planted in blocks of 18 rows for a total area of 3600 m^2^. Rows were separated by approximately ∼45cm and the final distance between plants within rows was 0.18m. At both locations, there was no separation between rows planted to the four cultivars, but the experimental area was surrounded by a border of sugar beets to minimize edge effects. In some cases, the border rows included one or more of the sugar beet hybrids used in the main experiment. Sugar beets were harvested in early November. No fungicides were applied and the CLS infections were natural.

**Table 1.**
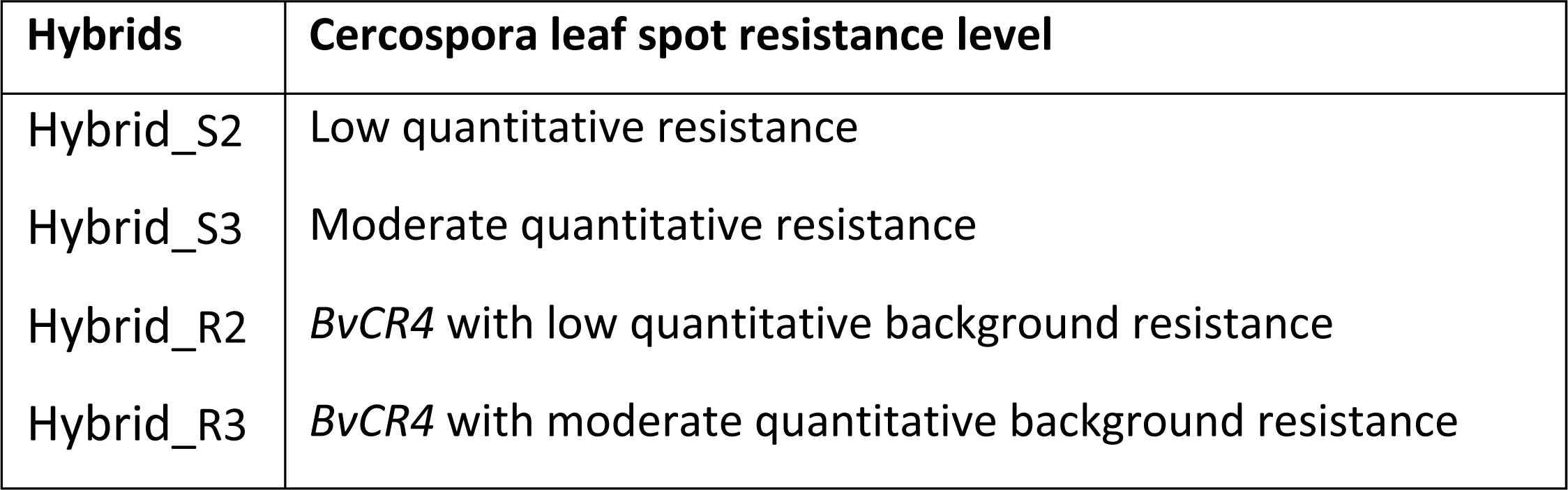
Sugar beet hybrids and their resistance to Cercospora leaf spot.

| Hybrids | Cercospora leaf spot resistance level |
| --- | --- |
| Hybrid_S2 | Low quantitative resistance |
| Hybrid_S3 | Moderate quantitative resistance |
| Hybrid_R2 | *BvCR4* with low quantitative background resistance |
| Hybrid_R3 | *BvCR4* with moderate quantitative background resistance |

### Temporal sampling strategy

Our sampling was designed to identify the naturally infected plants in each plot and determine how natural infection spreads among plants. We systematically searched each plot for infected plants once per week from July 21^st^ to September 24^th^, 2020. Individual sugar beet plants served as sampling units and one leaf with CLS lesions was collected from a plant. The sampling method is shown in Figure S1a. All sampled plants were marked with flags to prevent resampling. In the early stages of the epidemic, we found single newly infected plants surrounded by healthy plants, which we referred to as solitary infections. As the epidemic progressed, the disease often spread from a single infected plant to neighboring plants to form a cluster of infected plants, hereafter called a hotspot. For each solitary infection, we recorded the GPS location (Garmin) and gave it a two-letter code (e.g. AA) and the number 1 to indicate it was the first infected plant at that GPS position (e.g. AA1). If the infected plant was immediately adjacent to a previously identified and flag-labeled plant, it was considered part of the same hotspot. Infected plants in the same hotspots were given the same alphabetic code and a number to indicate the sampled plants were the 2nd, 3rd, 4th or 5th plant found in the hotspot (e.g. AA4 was the fourth plant sampled in the AA hotspot). As mentioned, one leaf with CLS lesions was taken from each plant, with a preference for young and otherwise healthy leaves with multiple clear CLS lesions. The excised leaves were kept flat and cool and transported back to the laboratory for *C. beticola* isolation (see below for details). We sampled a maximum of five plants from each hotspot, which typically covered an area of ∼1m^2^ when sampling ceased. We stopped sampling a hybrid plot when it was no longer possible to find new solitary infected plants because hotspots had coalesced. Our sampling design allowed us to track CLS epidemic progression and track hotspot formation within and between plots planted to resistant and susceptible hybrids.

### Leaf storage

To store leaves flat, each leaf was placed on a labeled piece of paper and covered with paper towel. Multiple leaves were stacked between stiff cardboard sheets and secured with strong rubber bands. The stacked leaves were then stored in a cool box with ice blocks to maintain a low temperature during transport. Upon arrival at the laboratory, the leaves were air dried at room temperature in a drying cupboard. Dried leaves were stacked and placed in plastic bags with anhydrous silica gel and stored at 4°C until used for pathogen isolation.

### Pathogen isolation and DNA extraction

To obtain pure strains of *C. beticola*, spores were collected from the dried leaves. Three or four pieces of leaf tissue with visible lesions were cut from each dried leaf. The tissue samples were surface sterilized by immersing them in a 2% sodium hypochlorite solution for 3 minutes, followed by two rinses in sterile de-ionized water. The leaf pieces were placed on dry paper towels to remove excess water, then were put into glass Petri dishes on an elevated sterile net placed on a piece of moist filter paper. The Petri dishes were sealed with parafilm and incubated at 24°C for 48-72 hours to induce sporulation. The induced lesions were checked for the typical black pseudostromata and hyaline conidia under a dissecting microscope (Olympus Schweiz AG). The spores were picked from sporulating lesions using hand-made fine glass needles and transferred to fresh potato dextrose agar (PDA, Difco). The plates were cultured under constant darkness at 24°C for 7-10 days. After a characteristic *C. beticola* colony had formed, a 5-mm agar plug was excised from the edge of each colony and transferred to a new PDA plate for 14 days to form a pure culture. Any contaminated plates were discarded. For long-term storage of each pure culture, 4-6 agar plugs were cut from the edges of a 2-week-old pure culture and put into cryotubes in duplicate - with and without anhydrous silica gel, and stored at 4°C. For most leaves, between 1-3 colonies were isolated from different lesions, but for some leaves, more than 3 strains were isolated to investigate pathogen diversity within the same leaf. In total, 1251 *C. beticola* isolations were made. For practical reasons, DNA was not extracted from all isolates. Instead, we aimed for a single isolation per leaf and preference was given to isolates from hotpots, which were expected to be more virulent and capable of dispersing to nearby healthy plants.

To obtain fresh mycelia for DNA extraction, we transferred hyphae from plates to liquid culture by scraping small pieces of hyphae from the edge of each pure culture and placing them into 150 mL flasks containing 50 mL potato dextrose broth (PDB, Difco). The liquid cultures were incubated in the dark at 24°C on a shaker at 120 rpm for 10 days. The fungal tissue was harvested, washed twice with sterilized water, and the supernatant was discarded after centrifuging (Allegra X-12R, Beckman Coulter) at 3750 rpm for 10 mins. The pellet was then transferred to 2 mL microcentrifuge tubes and lyophilized overnight. The lyophilized pellets were ground into a fine powder using ∼15 glass beads in a grinding machine (Qiagen MM300 Retsch TissueLyser). A total of 560 DNA samples of isolates coming from separate leaves were extracted using an automated DNA extraction robot, the Kingfisher Flex (Thermo Fisher Scientific), using the recommended reagents and a protocol for fungal samples (LGC Genomics GmbH). The resulting DNA samples were quantified using a Nanodrop spectrophotometer (Thermo Fisher Scientific). We performed a polymerase chain reaction (PCR) using species-specific primers (Knight and Pethybridge, 2020) to confirm the samples were *C. beticola*. Only samples with a clear band at around 200 bp (∼85%) were retained.

### Simple sequence repeats analysis

The six SSRs were amplified in 2 multiplexes (multiplex1: SSRCb1, SSRCb3, SSRCb21; multiplex2: SSRCb22, SSRCb25, SSRCb27) (Groenewald et al., 2007; Vaghefi et al., 2017b). The SSR primers were blasted against the reference genome to ensure they were not physically linked (Table S1). The PCR amplifications were performed using Qiagen Type-it kits (Qiagen) in a total volume of 11 μL. The reaction mixture contained 20 ng of template DNA, 0.4 μM of each forward and reverse primer, 4 μL Type-it mix (with buffer and MgCl_2_), 1 μL Q-solution (1 U Taq polymerase), and 2 μl deionized water. The initial denaturation was carried out at 95°C for 5 mins, followed by 30 cycles of denaturation at 95°C for 30 s, annealing at 58°C or 56°C for 90 s, elongation at 72°C for 30 s and final extension at 60°C for 30 mins. The PCR amplicons were analyzed on an ABI 3730xl DNA Analyzer (Applied Biosystems) at the Genetic Diversity Centre (GDC) of ETH Zürich. Multiple samples were repeated at different stages to create technical replicates to estimate SSR genotyping error rates and ensure consistent allele calling.

SSR allele size was scored using the R package *Fragman* (Covarrubias-Pazaran et al., 2016). To reduce the genotyping error, allele fragment sizes of each SSR marker were visualized and binned using the R package MsatAllele v 1.05 (Alberto, 2009). Fragment stutter patterns varied between the SSRs. Visual inspection of the allele calling and consistency between repeats was used to assess the accuracy of the MsatAllele automatic binning. Automatic binning was accurate for three SSRs (SSRCb27, SSRCb3, SSRCb21) but manual adjustment of the bin sizes was required for the rest. The consistency of allele calling was evaluated using isolates that were replicated (n=51). In total we obtained SSR genotypes for 475 isolates. Genotype calling was repeated for 20% of isolates chosen at random. Samples with multiple allele calls were assumed to be a mixture of strains and were excluded from the dataset. All individuals with missing SSR data were also removed, leaving 340 fully genotyped isolates (158 from Hendschiken and 182 from Rudolfingen) for downstream analyses.

### Statistical analysis

Statistical analysis was performed in R version 4.3.1 (R Core Team, 2023) and R Studio version 2023.6.0.421 (Posit team, 2023). To investigate the localized spread between plants to form a hotspot, we used a generalized linear model with a binomial distribution. To compare across hybrids, we needed to control for the fact that CLS continued to spread after we stopped sampling in some hybrid plots, meaning that any new infection cases that were identified at the end of the sampling period were not followed to determine if they spread to neighboring plants. Our solution was to right-censor the data by excluding the new infection cases observed 1 and 2 weeks before the final sampling date in each hybrid plot. This is a common practice when sampling ends before an event has been observed. As the sampling period varied for each hybrid, the right-censor date also varied (Table 2). The rate at which a hotspot developed was analyzed using a linear model. The pairwise distances between geotagged infection sites were calculated using the R package *geosphere* v1.5-18 (Hijmans, R. J., 2022). Whether the distributions of geotagged plants differed between hybrids or between the beginning and the end of the season were assessed using Kolmogorov-Smirnov (K-S) tests in the R package R4PDE (Del Ponte, 2023). Multi-locus genotypes (MLGs) were identified and analyzed with the R package *poppr* v.2.9.3 (Kamvar et al., 2014; Kamvar et al., 2015).

**Table 2.** The first appearance of CLS, the number of plants sampled at the first sampling date (No. plants), the number of days after the first appearance of symptoms on July 21^st^ (onset delay), the total length of the sampling period, and the last sampling date (the right censored date is shown in parenthesis, see Methods) for each hybrid at each field site.

| Location | Hybrid | First infection date | Onset delay | No. plants | Sampling period | Last sampling date (right censored date) |
| --- | --- | --- | --- | --- | --- | --- |
| Hendschiken | Hybrid_S2 | 07-21-20 | 0 | 22 | 30 | 08-20-20 (08-11-20) |
|  | Hybrid_S3 | 07-21-20 | 6 | 1 | 24 | 08-20-20 (08-11-20) |
|  | Hybrid_R2 | 08-11-20 | 21 | 6 | 33 | 09-08-20 (09-03-20) |
|  | Hybrid_R3 | 08-20-20 | 30 | 3 | 35 | 09-24-20 (09-03-20) |
| Rudolfingen | Hybrid_S2 | 07-21-20 | 0 | 7 | 30 | 08-20-20 (08-11-20) |
|  | Hybrid_S3 | 07-27-20 | 6 | 2 | 24 | 08-20-20 (08-11-20) |
|  | Hybrid_R2 | 08-25-20 | 35 | 43 | 14 | 09-08-20 (09-03-20) |
|  | Hybrid_R3 | 08-20-20 | 30 | 4 | 35 | 09-24-20 (09-03-20) |

We performed clone-correction for each site separately as our earlier analyses indicated it was unlikely to observe the same MLG between two sites (whole genome sequence data of 256 isolates found only one case of the same clone present at the two sites (Chen et al., 2024)). The number of alleles, number of unique MLGs, the expected number of MLGs controlling for differences in sample size (eMLG) and Nei’s gene diversity, Simpson’s diversity and evenness were calculated using the R package *poppr* v.2.9.3 for each hybrid and site. Bootstrapped confidence intervals were calculated for genetic diversity statistics using function diversity_ci in *poppr*. Clonal fraction was calculated as (N–number of MLGs)/N, where N equals the number of isolates.

## Results

### Onset and distribution of CLS disease

The first symptoms of *C. beticola* were observed in the susceptible hybrids: for Hybrid_S2 on July 21^st^ and one week later in Hybrid_S3. The first infections on the resistant hybrids appeared 21-30 days after the first infections appeared on the susceptible hybrids (Table 2). The appearance of the first *C. beticola* infections on resistant hybrids with *BvCR4* varied between fields. In Hendschiken, the disease appeared on Hybrid_R2 before Hybrid_R3, while in Rudolfingen, the first symptoms were observed later in Hybrid_R2. GPS coordinates were used to map the locations of infected plants. The spatial distribution of all infection cases is shown in Figure 1.

**Figure 1.**
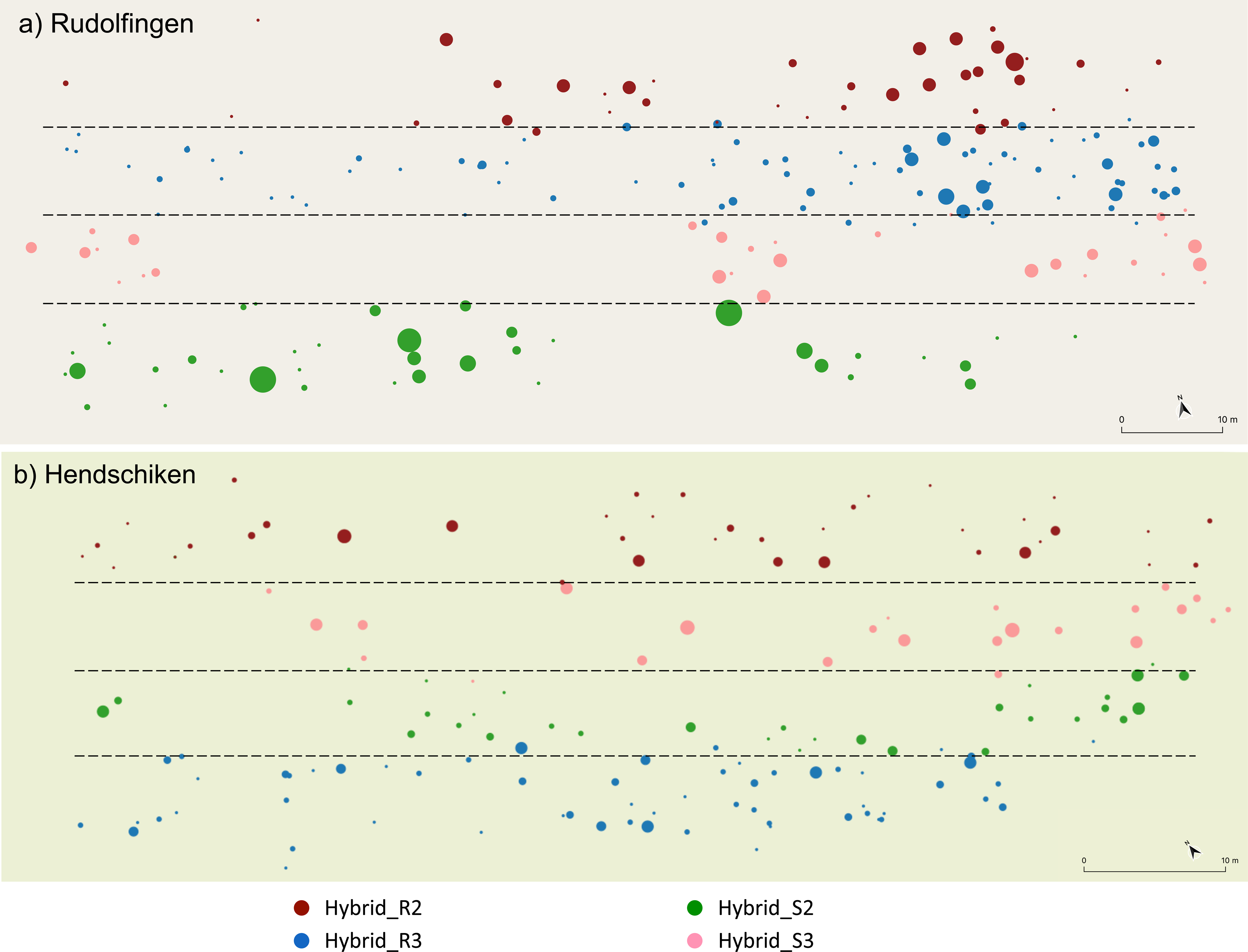
Spatial distribution of CLS in Rudolfingen (a) and Hendschiken (b). Each point represents a geotagged infection case. The size of the point indicates the number of sampled plants within the hotspot. The point color indicates the plant hybrid. Plant hybrid was determined by counting the planted rows from a reference point and using the different hybrid morphology. The dashed lines approximate the separation of the different hybrids, but there was no physical separation between hybrids. Thus, it can appear that one plot contains more than one hybrid (points with different colors), but this is an artifact of plotting and scaling the figure.

The accumulation of geotagged infection cases is shown in Figure 2. The sampling period varied for each hybrid because the dates of disease onset differed (Table 2) and because we stopped sampling when the hotspots coalesced and solitary infections could no longer be clearly distinguished. Thus, we stopped sampling first in Hybrid_S2, then Hybrid_S3 and then Hybrid_R2. Sampling continued until our final visit (September 24) in Hybrid_R3 as hotspots had not coalesced. Infections continued to increase in all hybrids after we stopped sampling. It should be noted that due to some initial sampling problems, we missed the start of the epidemic in Hybrid_S3. Hence caution must be taken when comparing the temporal spread in Hybrid_S3 with other hybrids. By the end of the season, 454 plants were sampled (one leaf per plant) from 196 geotagged infections in Hendschiken, and 503 plants were sampled from 209 geotagged infections in Rudolfingen.

**Figure 2.**
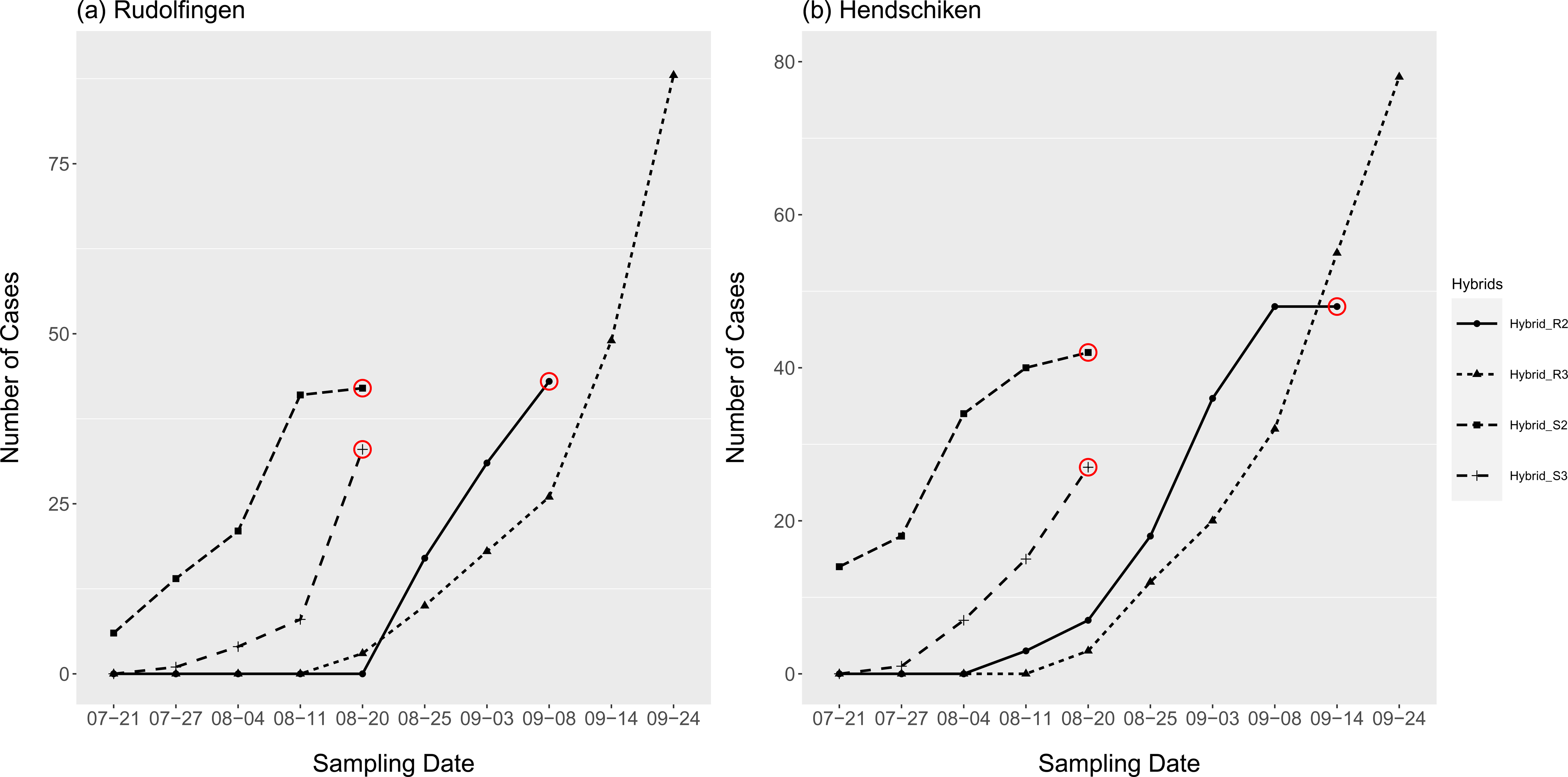
Accumulation of geotagged infection cases over time in Rudolfingen (a) and Hendschiken (b) in each hybrid. Sampling was terminated on different dates in each hybrid (indicated by red circles) when the disease had progressed to a point where new isolated infections could not be differentiated from already sampled hotspots (see methods and Table 2).

### Localized spread of CLS in each hybrid

Across all hybrids, about 80% of infection cases spread to at least one neighboring plant, resulting in the formation of hotspots (Table 3). There was no significant difference between fields or hybrids in the probability that CLS spread from one solitary infected plant to neighboring plants to form a hotspot of two or more infected plants, or to form a large hotspot with at least five infected plants (hotspot: Field *F*_(1,209)_ =1.23, *p*=0.26; Hybrid *F*_(3,206)_ =4.84, *p*=0.1, Large hotspot Field *F*_(1,209)_ =2.69, *p*=0.10; Hybrid *F*_(3,206)_ =4.90, *p*=0.17). NB: the numbers in Table 3 differ slightly compared to the total cases and hotspots (Figure 2) due to the right-censoring of datasets.

**Table 3.** The number of infection cases (IC) and the proportion of these that developed into hotspots (≥2 infected plants) and the proportion that developed into large hotspots (≥5 infected plants) for each hybrid. The data has been right censored (see Methods).

| Location | Hybrid | No. IC | Number of hotspots<br>(proportion) | Number of large hotspots<br>(proportion) |
| --- | --- | --- | --- | --- |
| Hendschiken | Hybrid _S2 | 37 | 26 (0.70) | 5 (0.14) |
|  | Hybrid _S3 | 20 | 17 (0.85) | 4 (0.20) |
|  | Hybrid _R2 | 22 | 18 (0.82) | 7 (0.32) |
|  | Hybrid _R3 | 30 | 24 (0.80) | 4 (0.13) |
| Rudolfingen | Hybrid _S2 | 23 | 19 (0.83) | 9 (0.39) |
|  | Hybrid _S3 | 18 | 17 (0.94) | 6 (0.33) |
|  | Hybrid _R2 | 29 | 26 (0.90) | 9 (0.31) |
|  | Hybrid _R3 | 31 | 23 (0.74) | 5 (0.16) |

To assess the rate of hotspot development in each hybrid, we calculated the rate of spread as the number of infected plants within a hotspot divided by the number of days between sampling periods for plants sampled in each hotspot. There was no difference in the rate of spread between sites (Field *F*_(1,115)_ =2.87, *p*=0.09), there was a significant effect of hybrid (Hybrid *F*_(3,115)_ =6.5, *p*<0.001) and no interaction (Field*Hybrid *F*_(3,115)_ =2.17, *p*=0.09). In Hendschiken, the rate of spread was slower in Hybrid_R3 and faster in Hybrid_R2 compared to Hybrid_S2 (Hybrid_R2: *t*=3.95, p=0.002, Hybrid_R3: *t*=-2.73, p=0.008). In Rudolfingen, there was no difference in the rate of spread, although Hybrid_R3 tended to be slower (*t*=-1.82, p=0.07).

### Spatial distribution of infections

We compared the distribution of geotagged infections at the beginning and the end of the epidemic and across hybrids within each site. The first two sampling dates in each hybrid plot were designated as the beginning of the epidemic and were used to investigate the spatial distribution of the first CLS infections. The initial distributions of hotspots and infection sites for most hybrids were randomly distributed (Table S2). During the epidemic, the minimum distance between infection sites decreased or remained constant, while the maximum distance increased throughout each epidemic (Table 4). The pairwise distances between hotspots ranged from ∼1 to 116 m over the course of the epidemics, with average values of 24 m and 36 m in Hendschiken and Rudolfingen, respectively. In some cases, the GPS was not recorded for an infection site, resulting in smaller numbers in Table 4 compared to Figure 2. The final distributions of hotspots and infection sites for most hybrids were aggregated (non-random) (Table S2) with two exceptions: Hybrid_R2 in Hendschiken and Hybrid_S2 in Rudolfingen, where the infection sites were randomly distributed. Although the number of infection cases increased over the season and were more likely to be aggregated, in pairwise comparisons of the spatial distribution of geotagged infections at the beginning and at the end of the season few significant differences were found (Table S3, Figure S2).

**Table 4.** The number of geotagged infection sites and the minimum, mean and maximum pairwise distance (m) between infection sites at the beginning and the end of the sampling period in each hybrid.

| Location | Hybrid | Pairwise distance at the start of epidemics |  |  |  | Pairwise distance at the end of epidemics |  |  |  |
| --- | --- | --- | --- | --- | --- | --- | --- | --- | --- |
|  |  | Count | Min | Mean | Max | Count | Min | Mean | Max |
| Rudolfingen | Hybrid _S2 | 14 | 2.2 | 28.2 | 73.7 | 41 | 1.3 | 36.7 | 110.7 |
|  | Hybrid _S3 | 4 | 3.8 | 54.7 | 98.6 | 31 | 1.3 | 47.8 | 116.3 |
|  | Hybrid _R2 | 15 | 1.3 | 19.7 | 62.0 | 37 | 1.3 | 29.2 | 108.3 |
|  | Hybrid _R3 | 8 | 4.2 | 33.9 | 94.5 | 88 | 0.2 | 37.2 | 110.0 |
| Hendschiken | Hybrid _S2 | 14 | 1.3 | 24.7 | 72.8 | 37 | 0.8 | 26.0 | 72.8 |
|  | Hybrid _S3 | 8 | 1.3 | 20.0 | 51.1 | 24 | 1.3 | 23.2 | 64.6 |
|  | Hybrid _R2 | 6 | 4.2 | 29.7 | 57.3 | 41 | 1.3 | 27.2 | 76.0 |
|  | Hybrid _R3 | 9 | 3.1 | 22.5 | 58.3 | 66 | 0.2 | 21.6 | 68.5 |

We compared the spatial distributions of geotagged infection sites in each hybrid and found that in Hendschiken Hybrid_R3 more often exhibited a different distribution of diseased plants compared to other hybrids (Table S4). In Rudolfingen, there were more differences: Hybrid_R2 was different from both susceptible hybrids, Hybrid_R3 was also different from Hybrid_S3, but was similar to Hybrid_S2, while Hybrid_S2 and Hybrid_S3 had different spatial distributions.

### Genetic and genotypic diversity at different hierarchical levels

The six SSR loci used in our study were polymorphic, resulting in 2 to 9 alleles per locus. Allelic diversity azer clone-correction ranged from 0.23 to 0.83 (Table S5) and there was some variation in the number of alleles for each SSR between hybrids (Table S6). There were some alleles unique to each field; for example, the 183 bp allele at SSRCb22 was found only in Hendschiken and the 373 bp allele at SSRCb27 was only detected in Rudolfingen.

We investigated genetic diversity at different hierarchical levels, including within and between hybrids and fields, and within leaves and hotspots. In Hendschiken, 158 isolates were distributed among 82 unique MLGs. In Rudolfingen, 182 isolates resulted in 77 unique MLGs (Table 5a). The genetic diversity ranged from 0.40 to 0.57 across different hybrid plots, and the genotypic diversity ranged from 0.73 to 0.95 (Table 5a). There was li}le difference in Nei’s gene diversity between sites or hybrids. The clonal fraction differed significantly among the hybrids in Rudolfingen (ξ^2^ = 11.15, df=3, p=0.01), while it was similar among hybrids at Hendschiken (χ^2^= 3.38, df=3, p=0.33). In Rudolfingen, the clonal fraction was lower in Hybrid_S3 and higher in Hybrid_R2. When resistant and susceptible hybrids are combined we saw a significantly higher clone fraction on resistant hybrids in Rudolfingen but not in Hendshicken (Table 5b). The frequency of MLGs differed in each site (Figure 3). In Rudolfingen, there was greater skew in MLG frequencies; R_MLG77 appeared in 46 isolates (20 in Hybrid_R2 and 26 in Hybrid_R3) and was restricted to the resistant hybrids with *BvCR4*. In Hendschiken, differences between the hybrids were less pronounced, with the most frequent MLG (H_MLG75) appearing 13 times in the resistant hybrids with *BvCR4* (Hybrid_R2 and Hybrid_R3) (3 and 10 times respectively) (Figure 3). The evenness was significantly higher in susceptible hybrids in Rudolfingen, but no difference was found in Hendschiken (Figure 4).

**Figure 3.**
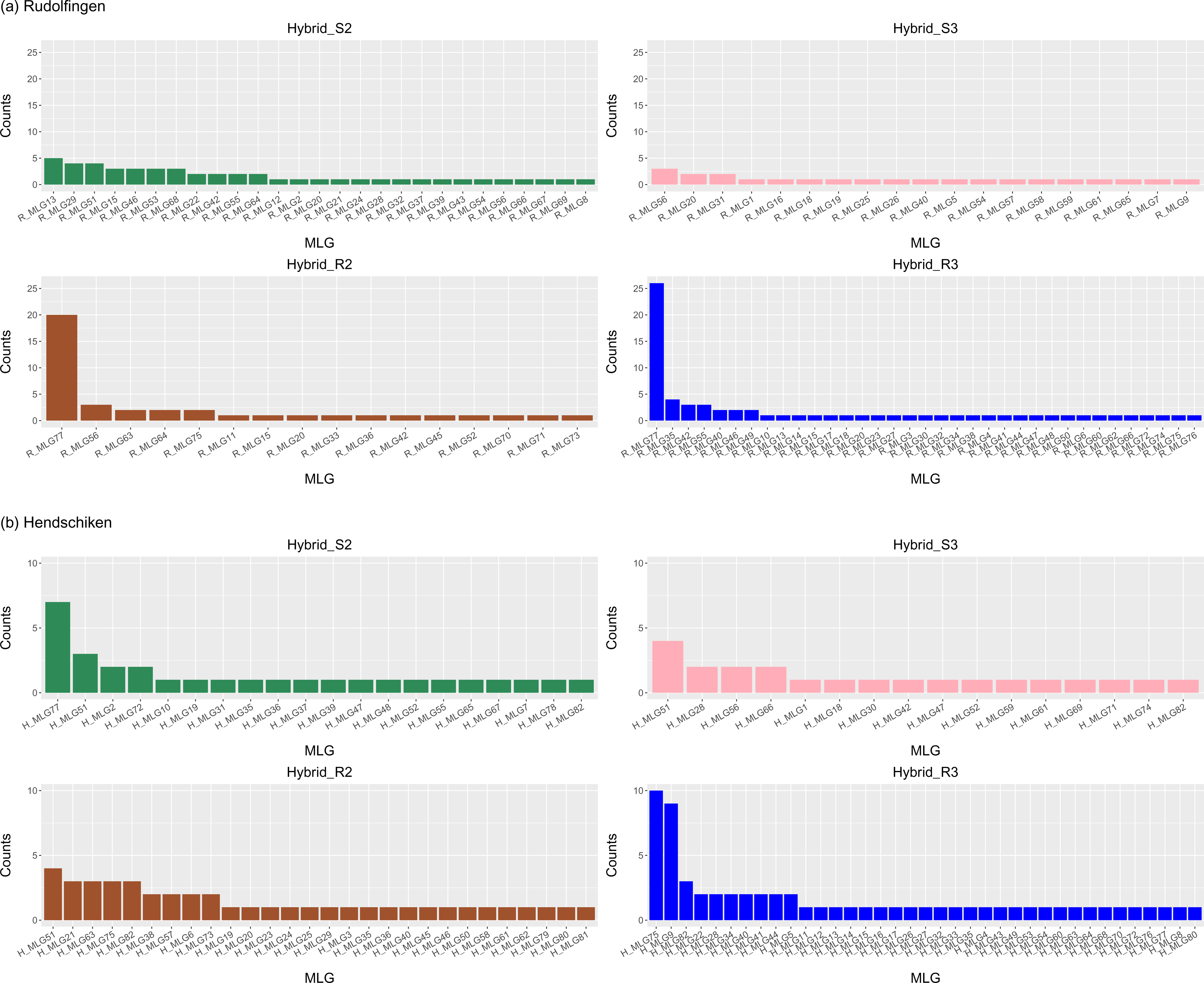
Frequency of each multilocus genotypes (MLGs) in each hybrid in Rudolfingen (a) and Hendschiken (b).

**Figure 4.**
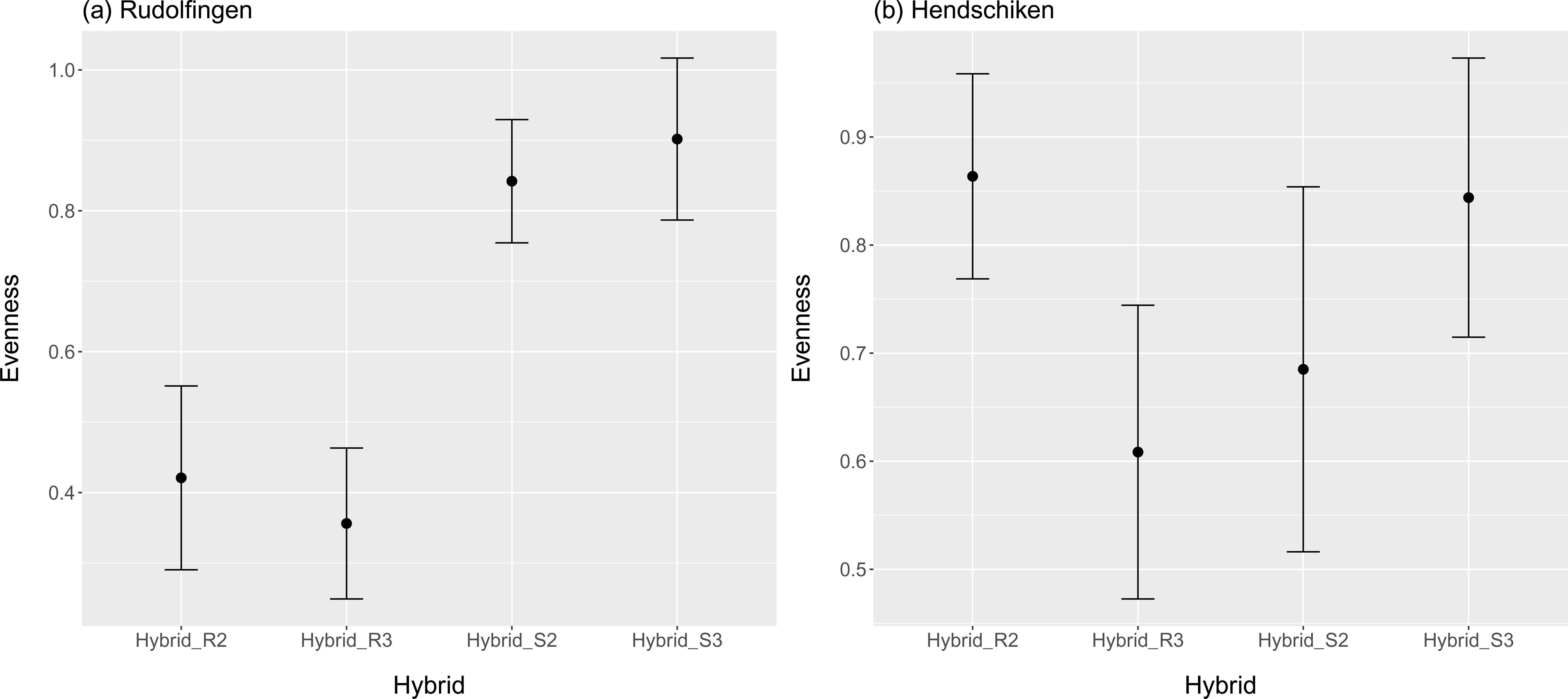
Evenness of multilocus genotypes (MLGs) in each hybrid in Rudolfingen (a) and Hendschiken (b). Points are the values of evenness and the error bars indicate the bootstrapped 95% confidence intervals.

**Table 5.** Indices of gene and genotype diversity based on six SSR loci for *Cercospora beticola* populations collected from a) four sugar beet hybrids in two fields in Switzerland. Table includes the number of strains (N), the number of multilocus genotypes (MLGs), the expected number of MLGs after rarefaction (eMLGs), the clonal fraction (CF=1-(MLGs/N), the Simpson’s complement index (λ) and Nei’s index of gene diversity before clone correction (He_0_) and after clone correction (He_cc_). b) Genotype diversity compared between Susceptible (BvCR-) and Resistant (BvCR+) hybrids combibned.

| Location | Hybrid | N | MLGs | eMLGs | CF | $\lambda$ | $H_{e0}$ | $H_{ecc}$ |
| --- | --- | --- | --- | --- | --- | --- | --- | --- |
| **Hendschiken** | Hybrid_S2 | 30 | 20 | 15.6 | 0.33 | 0.91 | 0.56 | 0.60 |
|  | Hybrid_S3 | 22 | 16 | 16 | 0.27 | 0.92 | 0.51 | 0.54 |
|  | Hybrid_R2 | 43 | 28 | 17.3 | 0.35 | 0.95 | 0.52 | 0.54 |
|  | Hybrid_R3 | 63 | 37 | 16.2 | 0.41 | 0.94 | 0.56 | 0.58 |
|  | **Total/Mean** | **158** | **82** | - | **0.48** | **0.97** | **0.56** | **0.56** |
| **Rudolfingen** | Hybrid_S2 | 49 | 27 | 16.7 | 0.45 | 0.95 | 0.51 | 0.54 |
|  | Hybrid_S3 | 23 | 19 | 19 | 0.17 | 0.94 | 0.57 | 0.60 |
|  | Hybrid_R2 | 40 | 16 | 10.7 | 0.60 | 0.73 | 0.40 | 0.56 |
|  | Hybrid_R3 | 70 | 35 | 14.1 | 0.50 | 0.85 | 0.52 | 0.57 |
|  | **Total/Mean** | **182** | **77** | - | **0.58** | **0.92** | **0.55** | **0.57** |

b)
| Location | Hybrid | N | MLGs | CF | Chisq | p-value |
| --- | --- | --- | --- | --- | --- | --- |
| **Hendschiken** | Susceptible | 52 | 36 | 0.31 |  |  |
|  | Resistant | 106 | 65 | 0.39 | 0.63 | 0.42 |
| **Rudolfingen** | Susceptible | 72 | 46 | 0.36 |  |  |
|  | Resistant | 110 | 51 | 0.54 | **4.68** | **0.03** |

### Genotypic diversity within a leaf and hotspot

To investigate genotypic diversity within individual leaves, multiple isolations were made from the same leaf for a total of 48 leaves (Table 6a). The leaves included in this analysis came from all four hybrids, were sampled over the time course of the epidemic, and had typical levels of infection, with at least 10-15 discrete CLS lesions distributed across the leaf. It was common to find more than one genotype per leaf, with 23 out of the 48 (0.48) tested leaves having more than one genotype on the same leaf. The maximum number of MLGs from one leaf was two, irrespective of the number of isolations (2-4). Although there was not enough data to make statistical comparisons between hybrids or sites, we commonly observed the coexistence of multiple genotypes in resistant hybrids. Specifically, in Hendschiken 6 out of 14 (0.39) tested leaves from resistant hybrids had multiple strains and 10 out of 16 (0.63) resistant leaves had multiple strains in Rudolfingen. We also considered the genotypic diversity within a hotspot (n= 48) (Table 6b). Just over half of the hotspots had more than one MLG. In resistant hybrids, 8 out of 13 (0.61) tested hotspots had multiple strains in Hendschiken and 11 out of 19 (0.57) hotspots had multiple strains in Rudolfingen.

**Table 6.**
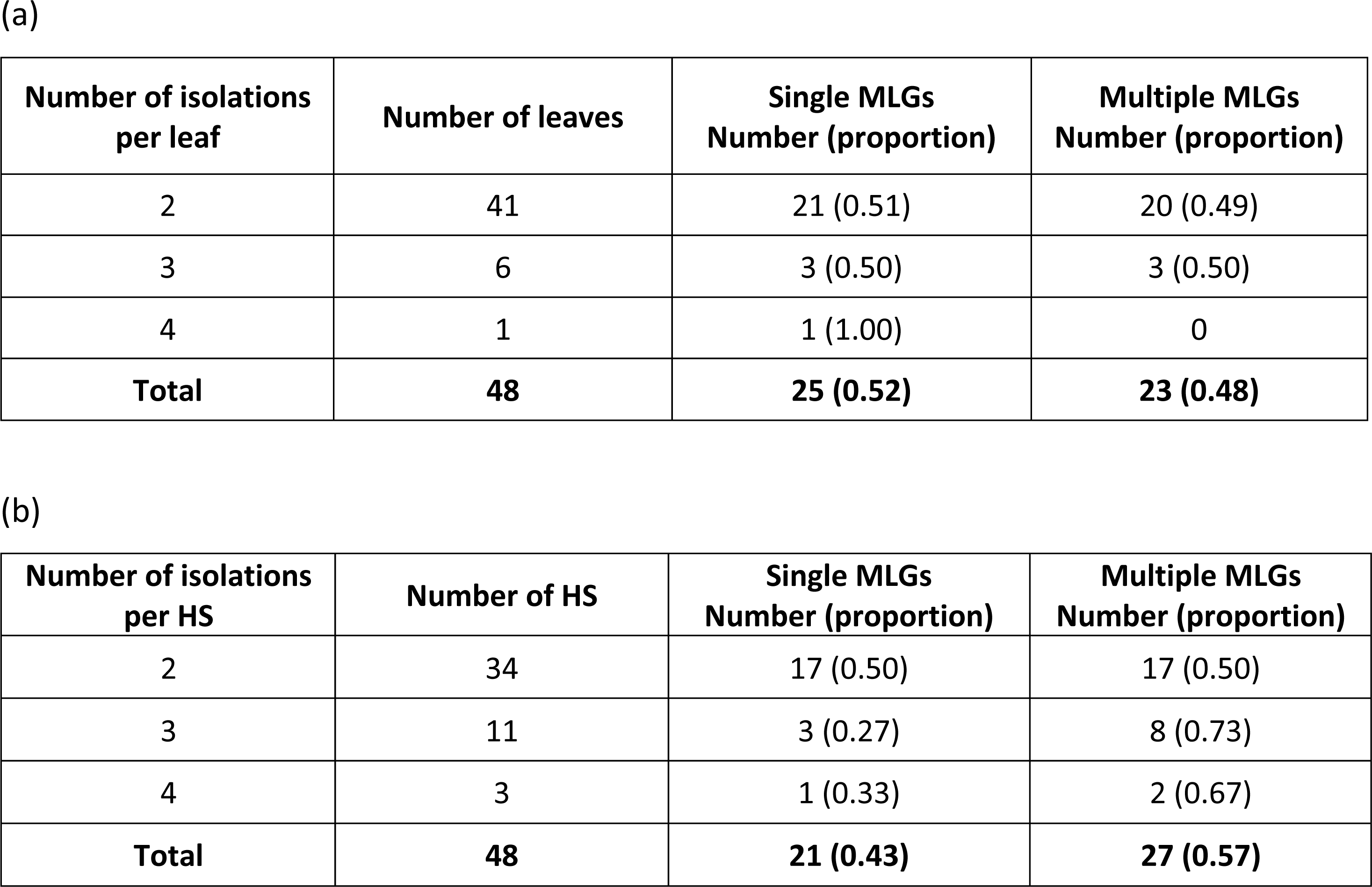
Genotypic diversity within (a) a leaf and (b) a hotspot (HS). The first column indicates a frequency class relating to the number of isolations per leaf/hotspot. There are three classes: 2, 3 or 4 isolations were made per leaf leaf/hotspot. The table includes the number of leaves/hotspots that had only one MLG (all clones) and with more than one MLG. The proportions are provided in parenthesis.

| Number of isolations per leaf | Number of leaves | Single MLGs<br>Number (proportion) | Multiple MLGs<br>Number (proportion) |
| --- | --- | --- | --- |
| 2 | 41 | 21 (0.51) | 20 (0.49) |
| 3 | 6 | 3 (0.50) | 3 (0.50) |
| 4 | 1 | 1 (1.00) | 0 |
| **Total** | **48** | **25 (0.52)** | **23 (0.48)** |

(b)
| Number of isolations per HS | Number of HS | Single MLGs<br>Number (proportion) | Multiple MLGs<br>Number (proportion) |
| --- | --- | --- | --- |
| 2 | 34 | 17 (0.50) | 17 (0.50) |
| 3 | 11 | 3 (0.27) | 8 (0.73) |
| 4 | 3 | 1 (0.33) | 2 (0.67) |
| **Total** | **48** | **21 (0.43)** | **27 (0.57)** |

### Genotypic diversity at the start of each epidemic

Genotypic diversity and the spatial distribution of isolates from the first two sampling dates in each plot were used to infer the most likely sources of primary inoculum in each epidemic. In Hybrid_S2 (July 21^st^ and 27^th^) in Rudolfingen we genotyped 13 isolates coming from seven hotspots and identified nine MLGs (clonal fraction (CF) = 0.31). In Hendschiken, we genotyped 14 isolates from six hotspots and identified seven MLGs (CF = 0.50), with five isolates from one of the hotspots having the same MLG. At the beginning of the epidemic, no two hotspots had the same MLG, indicating that different pathogen strains established each hotspot. Only four out of the seven initial hotspots in Rudolfingen were geotagged, with pairwise distances ranging from 18.4 to 64.7 meters. Three out of the six initial hotspots in Hendschiken were geotagged, with pairwise distances ranging from 25.0 to 68.7 meters. For the first two sampling dates in Hybrid_S3, only one isolate was genotyped from each site, and thus the distribution of MLGs cannot be compared.

For the resistant Hybrid_R2: In Rudolfingen, on the first sampling date (August 25^th^) 15 isolates from 11 hotspots were genotyped. These contained six MLGs (CF = 0.6) with pairwise distances ranging from 1.3 to 49.2 m. At the beginning of this epidemic the R_MLG77 genotype was found 10 times in eight hotspots that were separated by pairwise distances ranging from 1.3 to 19.6 m. In Hendschiken, during the first sampling dates in Hybrid_R2 (August 11^th^ and 20^th^), five isolates were sampled from four hotspots composed of four MLGs (CF = 0.2), with the pairwise distances ranging from 4.2 to 61.4 m. No two hotspots shared a MLG.

For the resistant Hybrid_R3: In Rudolfingen, on the first two sampling dates (August 20^th^ and 25^th^), four isolates from four hotspots were genotyped and each had a unique MLG (CF = 0), with pairwise distances ranging from 5.7 to 46.0 m. The frequent MLG R_MLG77 appeared in one of these hotspots. In Hendschiken, 15 isolates sampled from eight hotspots belonged to nine MLGs (CF = 0.4) (four isolates from the same hotspot had the same MLG), with pairwise distances between hotspots ranging from 6.2 to 58.3 m. One MLG was shared between two hotspots that were 7.8 m apart.

### Distribution of isolates with the same MLGs over time

The primary CLS infections in each plot gave rise to asexual conidia that provided secondary inoculum. The spatial distribution of the resulting clones with identical MLGs provides insights into the dispersal of secondary inoculum during a growing season. Common MLGs, observed ≥ 5 times in each field (eight in Rudolfingen and five in Hendschiken), were used to calculate the dispersal distance of the same genotype (Table 7). The minimum pairwise distance of 0 m was not included in the table. It was common to find the same MLG separated by distances exceeding 10 meters. The maximum distance between isolates sharing the same MLG was 108.5 m in Rudolfingen and 68.7 m in Hendschiken.

**Table 7.** The median, mean and the maximum pairwise distance (m) between the most frequent MLGs that were found at least five times in a field site.

|  | MLG | Count | Median | Mean | Maximum |
| --- | --- | --- | --- | --- | --- |
| **Rudolfingen** | R_MLG77 | 46 | 16.6 | 22.8 | 108.5 |
|  | R_MLG56 | 7 | 43.6 | 38.9 | 100.5 |
|  | R_MLG13 | 6 | 7.5 | 16.3 | 49.8 |
|  | R_MLG42 | 6 | 58.4 | 48.0 | 107.8 |
|  | R_MLG15 | 5 | 51.9 | 43.5 | 105.0 |
|  | R_MLG20 | 5 | 14.9 | 28.5 | 74.0 |
|  | R_MLG46 | 5 | 31.9 | 23.2 | 64.4 |
|  | R_MLG55 | 5 | 19.2 | 30.0 | 77.1 |
| **Hendschiken** | H_MLG75 | 13 | 4.2 | 7.8 | 20.5 |
|  | H_MLG51 | 11 | 15.7 | 18.9 | 46.7 |
|  | H_MLG9 | 9 | 3.7 | 4.2 | 9.4 |
|  | H_MLG77 | 8 | 15.0 | 13.5 | 36.7 |
|  | H_MLG82 | 8 | 29.3 | 27.1 | 68.7 |

## Discussion

The major objective of this study was to investigate how host resistance influenced CLS epidemics on sugar beet. We combined detailed epidemiological surveys with geo-referenced sampling and SSR genetic data to understand the fine scale temporal and spatial distribution of *C. beticola* genotypes on different resistant hybrids grown under the same environmental conditions during a single season. The dynamics of different pathogen populations were assessed on their respective hybrids during each epidemic. Resistant hybrids delayed the onset of epidemics, however, the host genotype did not affect the likelihood of spread to neighboring plants. No differences in genetic diversity were found between pathogen populations on different hosts, but we found high frequencies of some MLGs on resistant hosts. The distribution of CLS hotspots tended to be random at the beginning and aggregated at the end of each epidemic in each hybrid, with a few exceptions. We found that a single pathogen genotype could disperse over a hundred meters during an epidemic, illustrating the possibility of long-distance dispersal of individual clones within fields during a single growing season.

### Onset of the disease varied according to host genotype

CLS emerged at least 3 weeks later in the hybrids with *BvCR4* (Hybrid_R2 and Hybrid_R3) compared to the hybrids without *BvCR4* (Hybrid_S2 and Hybrid_S3). The delayed appearance of CLS in resistant hybrids at both sites suggests that the resistant hybrids suppressed the establishment of CLS early in the season, agreeing with observations from earlier field trials conducted by KWS (personal communication). Similar delays (2 to 4 weeks) in the onset of CLS on other resistant sugar beet hybrids were also reported in previous studies (Karaoglanidis et al., 2000; Weltmeier et al., 2011; Wolf and Verreet, 2005). It is possible that we did not always detect the start of an epidemic due to the size of the plots and our sampling frequency. For example, for Hybrid_R2 in Rudolfingen and Hybrid_S2 in Hendschiken, we collected leaves from 43 and 22 plants respectively on the first day of sampling (Table 2), suggesting that the epidemic had already started in these plots in the days between our sampling visits. For the six remaining hybrid plots, the number of plants collected on the first sampling date was <10, suggesting that we caught the beginning of those epidemics. If we consider only the six plots with less than 10 infected plants at the onset of the epidemic, the average distance between infected plants was 23.0 m in Hendschiken and 30.8 m in Rudolfingen. If we add the other two plots, the average distance between infected plants was 23.9 m in Hendschiken and 25.7 m in Rudolfingen. Given that sugar beets had not been grown at either field site for at least three years, our interpretation in both cases is that the primary inoculum was airborne. This is consistent with previous work demonstrating experimentally that the inoculum is dispersed mainly by wind (Khan et al., 2008) and originates external to the field of interest (Knight et al., 2018). Given the high genetic diversity found in the pathogen populations on these first infected plants and the random distributions of primary infections across the fields in both locations (Table S2), we hypothesize that the primary inoculum was mainly airborne ascospores. Though the sexual stage of *C. beticola* has not yet been described, most population genetics studies conducted thus far have indicated that sexual recombination is likely occurring regularly in this pathogen (Birla et al., 2012; Groenewald et al., 2007b, 2006) and its closest relatives include Mycosphaerella teleomorphs that are known to produce airborne ascospores (Goodwin et al. 2001).

### Localized spread was not strongly affected by host genotype

Host genotype did not affect the probability that a single infected plant would develop into a hotspot, with about 80% of infection cases spreading to adjacent plants (Table 3), but the speed of localized spread (involving 2-5 plants) varied between hybrids and fields. In Rudolfingen, there was no difference among hybrids in how quickly CLS spread to neighboring plants, but in Hendschiken, the spread was slower in Hybrid_R3 and faster in Hybrid_R2 compared to the susceptible Hybrid_S2. This field-dependent pattern may be attributed to differences in the distribution of the hybrid plots, in particular the susceptibility of the hybrid planted in a neighboring plot. Previous studies suggested that the level of host resistance did not influence the latent period, however, susceptible hybrids produced more conidia per unit of necrotic leaf area (Oerke et al., 2019). Thus, plots planted next to susceptible hybrids may experience higher local disease pressure. In Hendschiken, Hybrid_R2 was planted next to the susceptible Hybrid_S3, while in Rudolfingen, Hybrid_R2 was planted next to the resistant Hybrid_R3, which might have acted as a buffer zone that reduced the influx of conidia onto Hybrid_R2 (Figure S1). In addition to differences in local spore production, differences in local microclimates may also have affected epidemic development at each field site (Imbusch et al., 2021; Rossi et al., 2000; Vereijssen et al., 2006). It also is possible that our weekly sampling interval was not frequent enough to accurately monitor localized spread, as some hotspots spread from one to five plants between two successive time points.

To summarize, our findings suggest that the host genotype strongly influenced the emergence of CLS but had little impact on the probability that CLS would spread to adjacent plants and form hotspots of infection. This is in line with previous work that demonstrated that the latency of spore production was not influenced by host resistance (Rossi et al., 2000). If *C. beticola* strains succeed in overcoming the resistance of a host and became established on a resistant plant, their ability to spread to neighboring plants does not differ from the speed of dispersal found on susceptible plants.

### Infection sites were not randomly distributed in fields

At the end of the season, CLS infection sites were mostly aggregated (not randomly distributed) with one exception (Table S2). The distributions of infections changed little between the onset and the end of the epidemic, except for Hybrid_S2 and Hybrid_R2 in Rudolfingen (Table S3). The minimum distance between infection cases was approximately one meter, while the maximum distance reached 116 meters (Table 4). In line with earlier findings, we observed qualitatively higher levels of CLS in Hybrid_S2, Hybrid_S3 and Hybrid_R2 compared to Hybrid_R3 at the end of the season in both sites. Lesions coalesced and caused wilting and death of entire leaves, leading to new growth from the roots in hybrids S2, S3 and R2. Hybrid_R3 did not exhibit obvious lesion coalescence, leaf death, or regrowth despite having many plants with CLS. This may reflect the shorter time interval between disease onset and final sampling in Hybrid_R3. Studies have shown that later infection results in lower yield losses for resistant varieties (Kaiser et al., 2010), and that the delayed CLS onset observed in resistant hybrids was positively correlated with sugar yield (Wolf and Verreet, 2009).

### Genotypic variation and spatial distribution of genotypes

Earlier studies illustrated how pathogen populations with higher genetic diversity can have a higher evolutionary potential to overcome R genes (McDonald and Linde, 2002; Stam and McDonald, 2018). In our study, the average Nei’s gene diversity based on six SSR markers ranged from 0.40 to 0.57 (Table 5a). This is similar to what has been found in sugar beet fields around the world (Knight et al., 2019). It is important to note that our estimates of genotypic diversity were limited because only six SSR loci were used to identify putative clones. To estimate how accurately the six SSRs identified clones, we compared the clones identified using the SSR results with clones identified using whole genome sequences for 113 isolates where we obtained both types of data (Chen et al., 2024). The six SSR loci correctly identified clones or unique genotypes in 84% of cases (47 out of 58 in Rudolfingen and 48 out of 55 in Hendschiken). The incorrect assignments were a mix of wrongly identifying two isolates as sharing the same genotype (underestimating genotypic diversity) and failing to identify two isolates as being clones (overestimating genotypic diversity) at roughly equal proportions (9.7% and 6.2% respectively). While this illustrates the potential errors of our SSR genotypes, we expect that this error rate will be randomly distributed across hybrids, so comparisons of relative genotypic diversity between hybrids remain valid.

The high genotypic diversity found in each population supports the hypothesis that *C. beticola* populations undergo regular cycles of sexual recombination, as proposed in earlier studies (Bolton et al., 2012; Groenewald et al., 2006; Groenewald et al., 2008; Vaghefi et al., 2017a). We detected high genotypic diversity at small spatial scales, with 23 out of 48 tested leaves infected by more than one MLG and 27 out of 48 tested hotspots containing more than one MLG. Other studies also observed multiple SSR MLGs on a single leaf in table beet and Swiss chard (Vaghefi et al., 2017c) and using other genetic markers (TAPD and DAMD) on sugar beet (Moretti et al., 2006). Our data suggests that co-infection is common on both resistant and susceptible hosts and that multiple strains may contribute to the spread of disease within a CLS hotspot.

We hypothesized that virulent isolates would be more prevalent on *BvCR4* hybrids due to their ability to overcome this resistance gene. If only a small fraction of isolates in the primary inoculum are virulent and able to produce asexual secondary inoculum on the resistant hybrids, we would expect these virulent clones to increase in frequency on resistant hybrids during the epidemic, resulting in lower genotypic diversity and higher clonal fractions on resistant hybrids compared to susceptible hybrids. Consistent with this expectation, we detected higher clone fraction in resistant (BvCR+) hybrids in Rudolfingen (Table 5b), but no difference at Hendschiken. Suggesting that the influence of this R gene on pathogen genotypic diversity can vary between locations. Consistent with the clone fraction variation, we observed skewed genotype frequencies on the resistant hybrids (Figure 3 and 4) most notably in Rudolfingen: one MLG (R_MLG77) appeared in 46 isolates exclusively in resistant Hybrid_R2 and Hybrid_R3. In Hendschiken, the differences in frequency were less striking, but one MLG (H_MLG75) appeared in 13 isolates solely found on the *BvCR4* hybrids. A subset of the isolates collected from resistant hybrids were tested for virulence against hybrids with and without *BvCR4* under field conditions in a later study. Under experimentally elevated disease pressure, that study confirmed that the highest frequency MLGs were virulent strains that could overcome *BvCR4* (Chen et al., 2024). It also found that only half of the isolates collected from hybrids with *BvCR4* were virulent against this gene. The lack of evidence for evolutionary bottlenecks in our study may be attributed to the frequent coexistence of virulent and avirulent strains within the same hybrid. Most of our temporal sampling occurred during the onset and development of the epidemic, thus we did not capture the end-of-season dynamics which might show stronger bottlenecks following clone competition. Transect or random sampling of the pathogen populations at the end of the season may provide a better test of the effectiveness of resistant hybrids as selection sieves.

### Source of inoculum and spatial distribution of MLGs

One of our aims in this study was to identify the potential sources of primary inoculum for CLS epidemics. We hypothesized that a CLS epidemic can be initiated by wind-dispersed ascospores which are capable of long-distance dispersal over hundreds of meters or by conidia on infected leaves from earlier crops that are moved by wind or watersplash. If the primary inoculum in a plot originates from windborne ascospores coming from outside the field, we would expect to find genetically different isolates sparsely and randomly distributed across individual plants at the onset of an epidemic. Alternatively, if the primary inoculum arises from locally produced asexual spores transmitted through watersplash from crop debris or human activity (e.g. infested seeds or machinery), we would expect to find the same genotype occurring simultaneously in different hotspots that might be aggregated in the field (for the case of infested machinery). In plots planted with Hybrid_S2, at both sites we observed unique genotypes in each hotspot, suggesting that unique genotypes led to the establishment of each hotspot, and that epidemics were initiated by genetically diverse primary inoculum coming from outside the fields. Similar patterns were observed in Hybrid_R3 at both sites and in Hybrid_R2 in Hendschiken, with no or few isolates shared between hotspots. But a different pattern was found for the resistant Hybrid_R2 in Rudolfingen. In this epidemic, the genotypic diversity of the first infections was lower, and the dominant genotype R_MLG77 appeared simultaneously 10 times in 8 hotspots distributed widely across the field. The most parsimonious explanation is that asexual spores originating from a neighboring unsampled border planted with the resistant hybrid may have disproportionally affected the genotypic diversity found in the sampled Hybrid_R2 plot. We postulate that the R_MLG77 strain emerged from standing genetic variation and attained a relatively high frequency in the border plots before being transmitted into the sampled Hybrid_R2 plot (Figure S1). Except for Hybrid_R2 in Rudolfingen, we found that genetically distinct primary inoculum was distributed widely across most plots (Table S2), as expected for wind-dispersed ascospores originating from outside of the field. This finding is in line with a previous study that found no sharing of MLGs between years, suggesting that the dominant source of inoculum for an epidemic is external to the field of interest (Knight et al., 2018).

Our experiment also aimed to better understand the dispersal pattern of secondary inoculum within a season. In Rudolfingen, high frequency MLGs (≥5) were separated by between 49 and 108 meters and in Hendschiken by between 9 and 68 m (Table 7). This is the first study in the CLS pathosystem to measure the dispersal of individual genotypes within a season. The dispersal distances we observe are unlikely to be due solely to rainsplash, as rainsplash can only disperse *C. beticola* spores short distances (several meters) (Carlson, 1967). Wind dispersal is a more likely mechanism, and although wind was already recognized as an important mechanism, it was not clear before how far wind could disperse *C. beticola* (Khan et al., 2008). Other possible dispersal mechanisms cannot be ruled out. It is possible that insects could also disperse the spores, along with human activity, including machine operations or spores moving on contaminated clothing during our weekly sampling visits.

## Acknowledgement

This project was funded by KWS SAAT SE & CO. KGaA. We are grateful to the KWS colleagues and farmers who contributed to the planting work. We thank Dr. Julien Alassimone, bachelor students Sophie Kuhn and Vivienne Hanke for assistance in the lab, and the Genetic Diversity Center (GDC) of Zurich for SSR fragment analysis.

## Data Availability Statement

The data that support the findings of this study are provided in the Supplementary information files.

