## Supplementary Table for "Molecular epidemiology of Cercospora leaf spot on resistant and susceptible sugar beet hybrids"

**Supplementary Tables and Figures**

**Table S1**. Information on the six polymorphic simple sequence repeats (SSR) used for genotyping, including the primer sequence, multiplex (MP) grouping, dye labelling, position on new reference genome (CB-Hev), GenBank accession and estimated error rate.

| SSR Name | Primer sequence | MP | Repeat motif | Dye | Regions on the chromosomes | GenBank Accession no. | Error rate |
|---|---|---|---|---|---|---|---|
| SSRCb22 | F: GCCACTTCATTACCACCTTGAAT<br>R: TGAGCTGATGTGAAAGGTAGAGG | 1 | (GAA) | Fam | Chr4:4057481-4057506 | KX452352 | 0.054 |
| SSRCb25 | F: GACGAGCATTCCATTGAGAAGTC<br>R: TCGTCGTTTTGGTCCTCTTCTTC | 1 | (GAG) | Hex | Chr10:1433597-1433622 | KX452355 | 0.054 |
| SSRCb27 | F: CGTCAAAGCAGTCCCTCGAT<br>R: AATTGAACAAGCGCCCAACC | 1 | (CAA) | Fam | Chr1:4165647-4165667 | KX452357 | 0.034 |
| SSRCb1 | F: TGCGATCTGGGCATAAATATC<br>R: AGATTTGCATTTGCCCACAC | 2 | (TC) | Hex | Chr2:329411-329433 | DQ902564 | 0.010 |
| SSRCb3 | F: ATAGAGTCAAACCAAGCCAAG<br>R: CCCGTTATAGCGCCCTTAG | 2 | (AC) | Fam | Ch9:1441450-1441472 | DQ902566 | 0.070 |
| SSRCb21 | F: GACTTTGGCATTCGAGAAGATGG<br>R: CCACTAAACGTATCTCTTTGCTGT | 2 | (AGC) | Fam | Chr2:2334406-2334431 | KX452351 | 0.052 |

**Table S2**. Kolmogorov-Smirnov (KS) tests of CLS spatial distribution in each hybrid. The test is performed by comparing the observed distribution with a random null distribution. Longitude was used as the spatial covariate and D represents the test statistic of the KS test.

| **KS test** | **Hendschiken** | | | | **Rudolfingen** | | | |
|---|---|---|---|---|---|---|---|---|
| | Initial | | All | | Initial | | All | |
| | D | *p*-value | D | *p*-value | D | *p*-value | D | *p*-value |
| **Hybrid_S2** | 0.36 | 0.07 | 0.35 | **<0.01** | 0.23 | 0.57 | 0.17 | 0.28 |
| **Hybrid_S3** | 0.27 | 0.29 | 0.30 | **0.04** | 0.34 | 0.42 | 0.26 | **0.03** |
| **Hybrid_R2** | 0.26 | 0.61 | 0.19 | 0.10 | 0.24 | 0.06 | 0.26 | **0.01** |
| **Hybrid_R3** | 0.31 | 0.30 | 0.18 | **0.02** | 0.34 | 0.18 | 0.22 | **<0.01** |

**Table S3**. Asymptotic pair-wise Kolmogorov-Smirnov (KS) tests between the initial and final CLS spatial distribution in each hybrid. D represents the test statistic of the KS test.

| **KS test** | **Hendschiken** | | **Rudolfingen** | |
|---|---|---|---|---|
| | **D** | ***p*-value** | **D** | ***p*-value** |
| **Hybrid_S2** | 0.03 | 0.97 | 0.18 | **0.01** |
| **Hybrid_S3** | 0.18 | 0.30 | 0.23 | 0.85 |
| **Hybrid_R2** | 0.15 | 0.88 | 0.22 | **<0.01** |
| **Hybrid_R3** | 0.07 | 0.99 | 0.18 | 0.30 |

**Table S4**. Asymptotic pair-wise Kolmogorov-Smirnov (KS) tests between the spatial distributions of all CLS infection sites in each hybrid at the end of the season. The null hypothesis is that the two compared distributions are the same.

| **KS test** | **Hendschiken** | | | | **Rudolfingen** | | | |
|---|---|---|---|---|---|---|---|---|
| **p-value / D** | **Hybrid_S2** | **Hybrid_S3** | **Hybrid_R2** | **Hybrid_R3** | **Hybrid_S2** | **Hybrid_S3** | **Hybrid_R2** | **Hybrid_R3** |
| **Hybrid_S2** | - | **0.03** | 0.10 | **<0.01** | - | **<0.01** | **<0.01** | 0.85 |
| **Hybrid_S3** | 0.11 | - | 0.13 | **0.07** | 0.20 | - | **<0.01** | **<0.01** |
| **Hybrid_R2** | 0.06 | 0.00 | - | **<0.01** | 0.13 | 0.28 | - | **<0.01** |
| **Hybrid_R3** | 0.15 | 0.18 | 0.14 | - | 0.02 | 0.19 | 0.14 | - |

**Table S5.** Summary of allelic diversity for each SSR locus at each field site, with the number of alleles (Na) and Nei's gene diversity (He) before ($He_0$) and after clone correction ($He_{CC}$).

| **Location** | **Hendschiken** | | | **Rudolfingen** | | |
|---|---|---|---|---|---|---|
| **Locus** | **Na** | **$He_0$** | **$He_{CC}$** | **Na** | **$He_0$** | **$He_{CC}$** |
| **SSRCb1** | 2 | 0.37 | 0.41 | 2 | 0.10 | 0.23 |
| **SSRCb21** | 6 | 0.81 | 0.82 | 6 | 0.78 | 0.83 |
| **SSRCb22** | 4 | 0.52 | 0.52 | 3 | 0.50 | 0.53 |
| **SSRCb25** | 4 | 0.66 | 0.68 | 4 | 0.66 | 0.64 |
| **SSRCb27** | 2 | 0.39 | 0.37 | 3 | 0.50 | 0.49 |
| **SSRCb3** | 8 | 0.63 | 0.65 | 9 | 0.75 | 0.76 |
| **mean** | **4.33** | **0.56** | **0.58** | **4.5** | **0.55** | **0.58** |

**Table S6.** Summary of allelic diversity in each Hybrid at (a) Hendschiken and (b) Rudolfingen. Each table includes the number of alleles ($N_a$) and Nei's genetic diversity (He) before ($He_0$) and after clone correction ($He_{CC}$).

(a) Hendschiken

| **Hybrid** | **Hybrid_S2** | | | **Hybrid_S3** | | | **Hybrid_R2** | | | **Hybrid_R3** | | |
|---|---|---|---|---|---|---|---|---|---|---|---|---|
| **Locus** | $N_a$ | **He** | $He_{CC}$ | $N_a$ | $He_0$ | $He_{CC}$ | $N_a$ | $He_0$ | $He_{CC}$ | $N_a$ | $He_0$ | $He_{CC}$ |
| **SSRCb1** | 2 | 0.29 | 0.34 | 2 | 0.17 | 0.23 | 2 | 0.34 | 0.35 | 2 | 0.46 | 0.45 |
| **SSRCb21** | 6 | 0.79 | 0.84 | 6 | 0.75 | 0.81 | 6 | 0.80 | 0.84 | 6 | 0.78 | 0.82 |
| **SSRCb22** | 4 | 0.58 | 0.62 | 2 | 0.45 | 0.46 | 2 | 0.49 | 0.52 | 2 | 0.51 | 0.51 |
| **SSRCb25** | 3 | 0.67 | 0.69 | 3 | 0.65 | 0.66 | 4 | 0.59 | 0.62 | 4 | 0.69 | 0.67 |
| **SSRCb27** | 2 | 0.46 | 0.48 | 2 | 0.52 | 0.50 | 2 | 0.34 | 0.30 | 2 | 0.40 | 0.35 |
| **SSRCb3** | 4 | 0.58 | 0.65 | 3 | 0.69 | 0.58 | 5 | 0.58 | 0.60 | 6 | 0.54 | 0.67 |
| **Total** | 21 | - | - | 18 | - | - | 21 | - | - | 22 | - | - |

(b) Rudolfingen

| **Hybrid** | **Hybrid_S2** | | | **Hybrid_S3** | | | **Hybrid_R2** | | | **Hybrid_R3** | | |
|---|---|---|---|---|---|---|---|---|---|---|---|---|
| **Locus** | $N_a$ | $He_0$ | $He_{CC}$ | $N_a$ | $He_0$ | $He_{CC}$ | $N_a$ | $He_0$ | $He_{CC}$ | $N_a$ | $He_0$ | $He_{CC}$ |
| **SSRCb1** | 2 | 0.08 | 0.14 | 2 | 0.30 | 0.35 | 1 | 0.00 | 0.00 | 2 | 0.11 | 0.21 |
| **SSRCb21** | 6 | 0.83 | 0.83 | 6 | 0.76 | 0.78 | 6 | 0.58 | 0.84 | 6 | 0.72 | 0.82 |
| **SSRCb22** | 2 | 0.38 | 0.43 | 3 | 0.56 | 0.57 | 3 | 0.27 | 0.54 | 2 | 0.40 | 0.51 |
| **SSRCb25** | 3 | 0.59 | 0.66 | 4 | 0.60 | 0.61 | 3 | 0.59 | 0.68 | 3 | 0.64 | 0.64 |
| **SSRCb27** | 2 | 0.50 | 0.48 | 3 | 0.53 | 0.53 | 2 | 0.33 | 0.53 | 2 | 0.50 | 0.48 |
| **SSRCb3** | 6 | 0.67 | 0.71 | 5 | 0.70 | 0.77 | 4 | 0.66 | 0.74 | 7 | 0.72 | 0.78 |
| **Total** | 21 | - | - | 23 | - | - | 19 | - | - | 22 | - | - |

**Figure S1**. A schematic of the sampling procedure and the arrangement of the hybrids plots at the two field sites.

**Figure S2**. Frequency distribution of pairwise distances between initial and final infection sites in each hybrid in Rudolfingen and Hendschiken. Initial: the first two sampling dates. Final: last sampling date. Different colors refer to different hybrids: green: Hybrid_S2; pink: Hybrid_S3; brown: Hybrid_R2; blue: Hybrid_R3.

**Figure S3.** Density plots of the pairwise distances between initial and final infection sites in each hybrid in Rudolfingen (left) and Hendschiken (right). Dashed lines indicate the initial sampling (the first two sampling dates), and the solid lines indicate the final sampling (last sampling dte). Different colors refer to different hybrids: green: Hybrid_S2; pink: Hybrid_S3; brown: Hybrid_R2; blue: Hybrid_R3.

**Figure S4.** Maximum distances between identical genotypes in fields, and the number of days between these observations in Rudolfingen (left) and Hendschiken (right).
