## Supplementary Figure for "Molecular epidemiology of Cercospora leaf spot on resistant and susceptible sugar beet hybrids"

**a) Near weekly temporal sampling.** We walked through the field site and sampled all diseased plants. We noted if a cluster of diseased plants (a hotspot) developed over time.

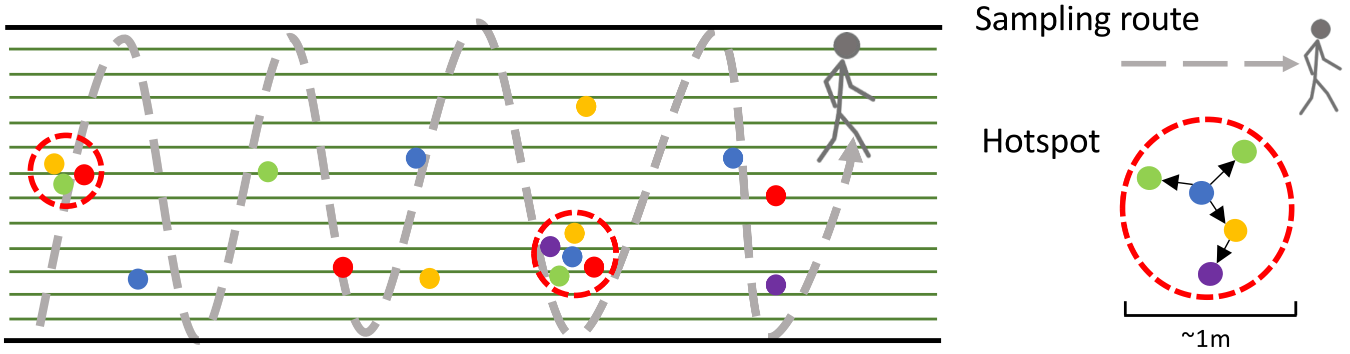


**b) The arrangement of sugar beet hybrids at the two field sites.** The experimental plots were sampled regions within a larger field planted with hybrids. The sampled area was surrounded by a buffer of at least 12 rows of sugar beet, to reduce edge effects. Each hybrid plot (12-18 rows) was directly adjacent to the neighbouring hybrid plot – there was no buffer between the hybrid rows.

**i) Hendschiken**

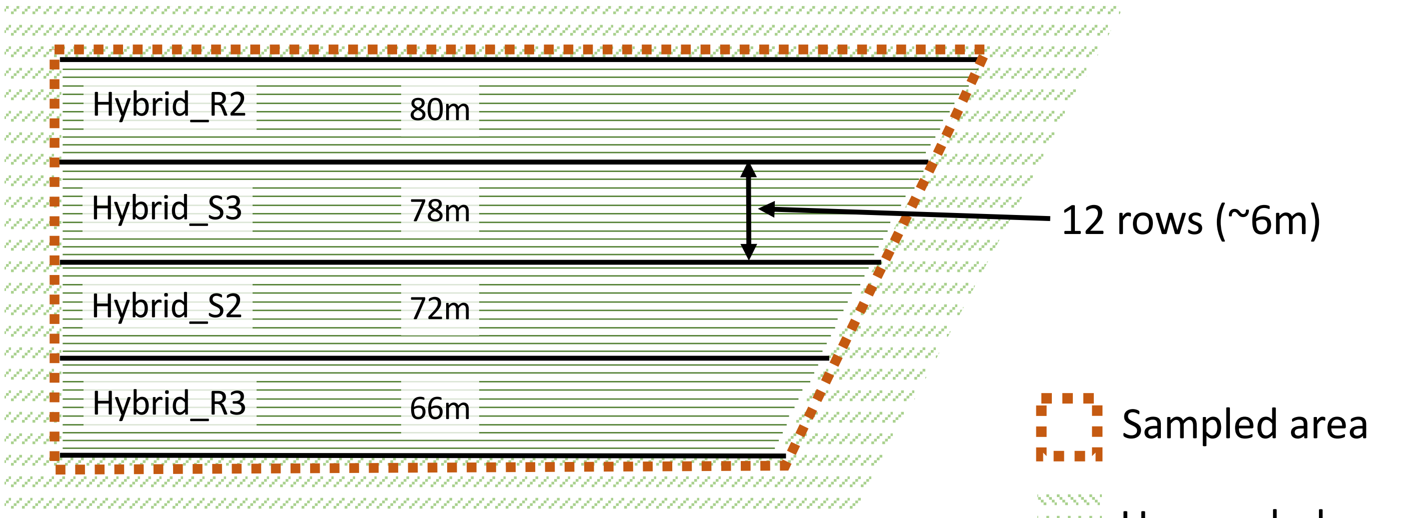


**ii) Rudolfingen**

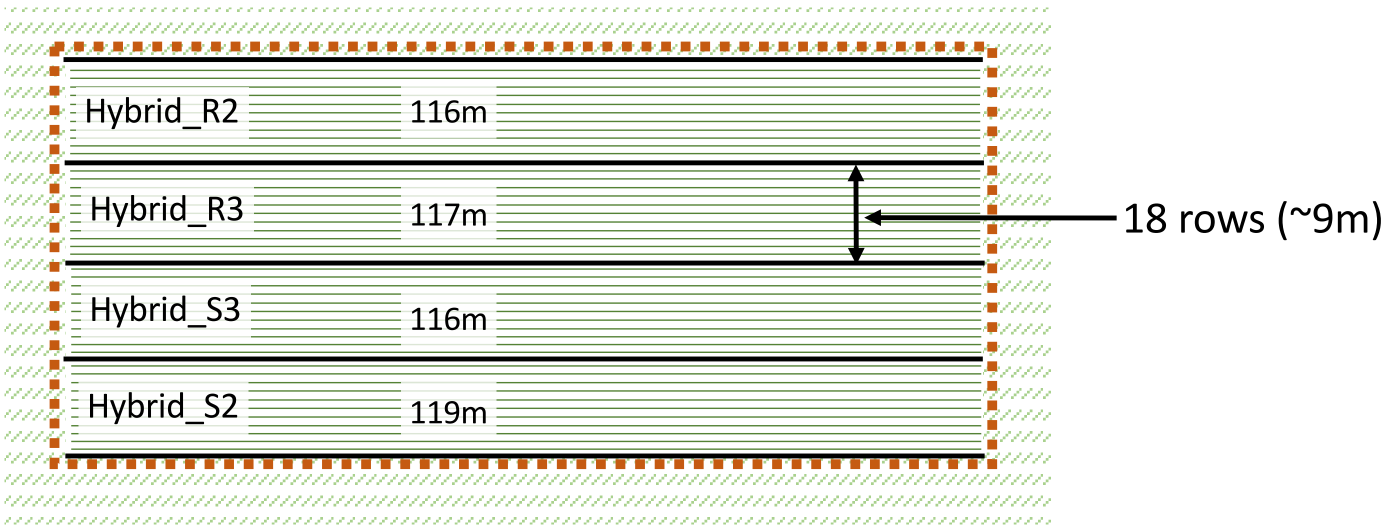

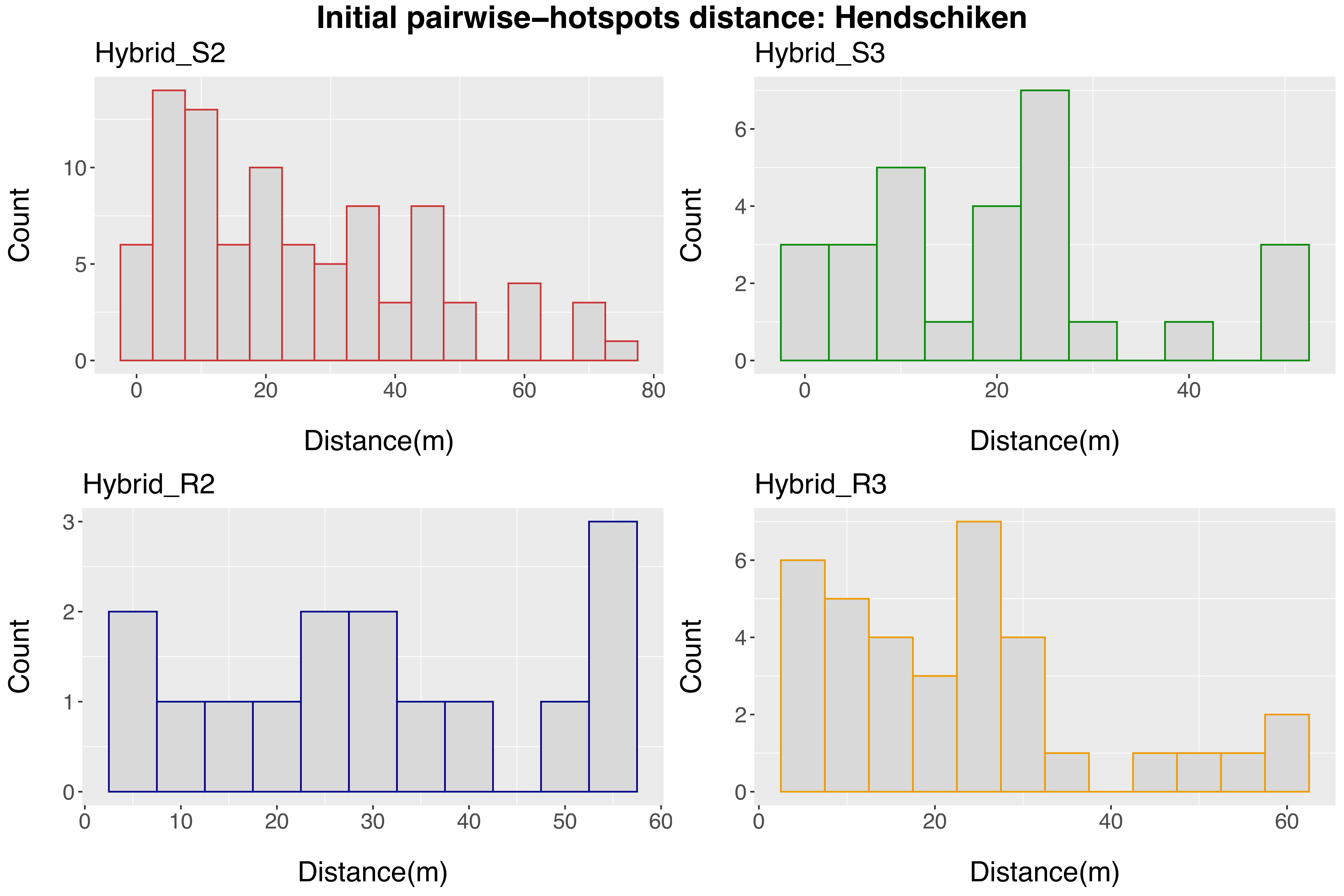

Initial pairwise−hotspots distance: Hendschiken
Hybrid_S2
Hybrid_S3
Hybrid_R2
Hybrid_R3
Count
Distance(m)
0
5
10
20
40
60
80
2
4
6
1
3
30
50

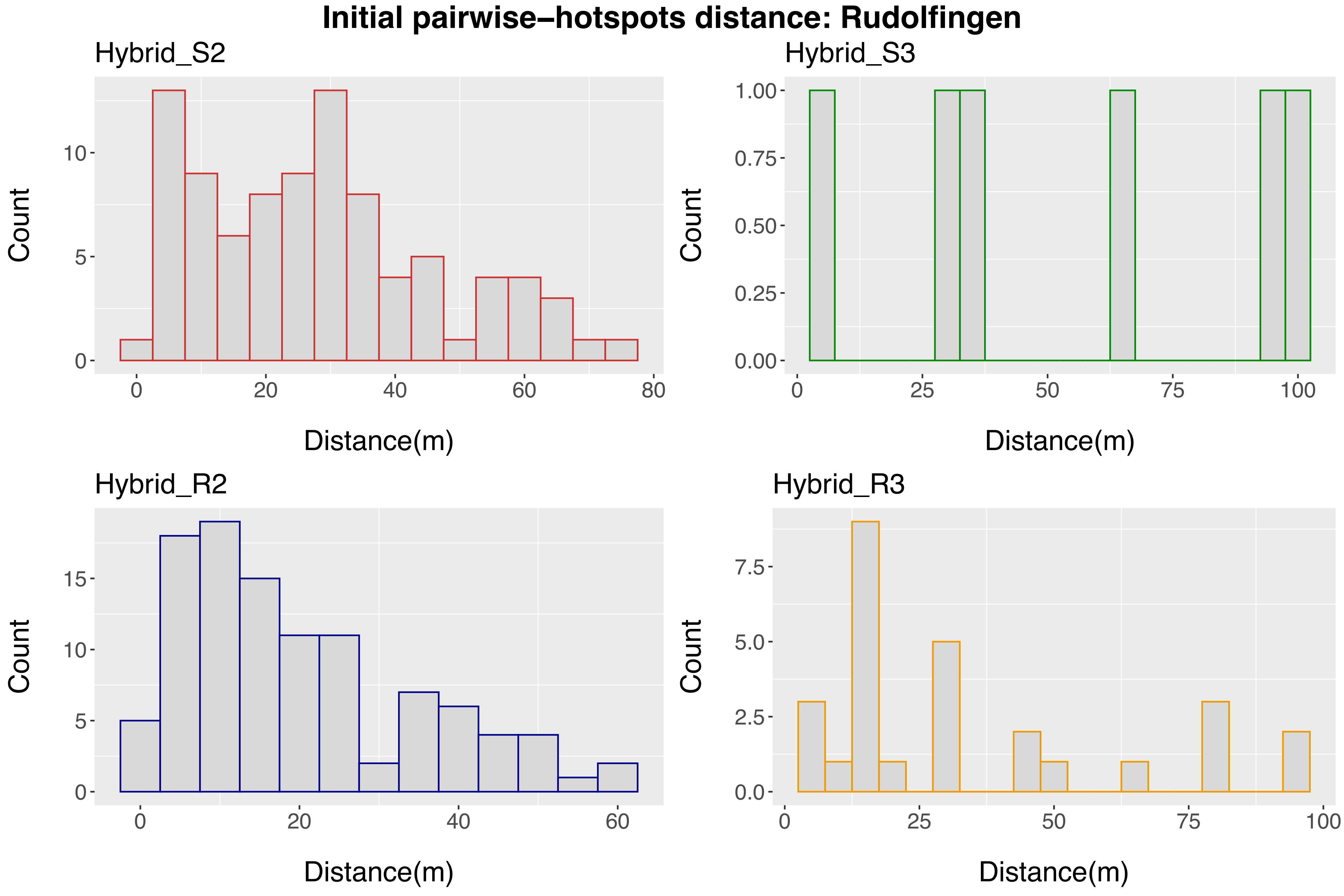

Initial pairwise–hotspots distance: Rudolfingen
Hybrid_S2
Hybrid_S3
Hybrid_R2
Hybrid_R3
Count
Distance(m)

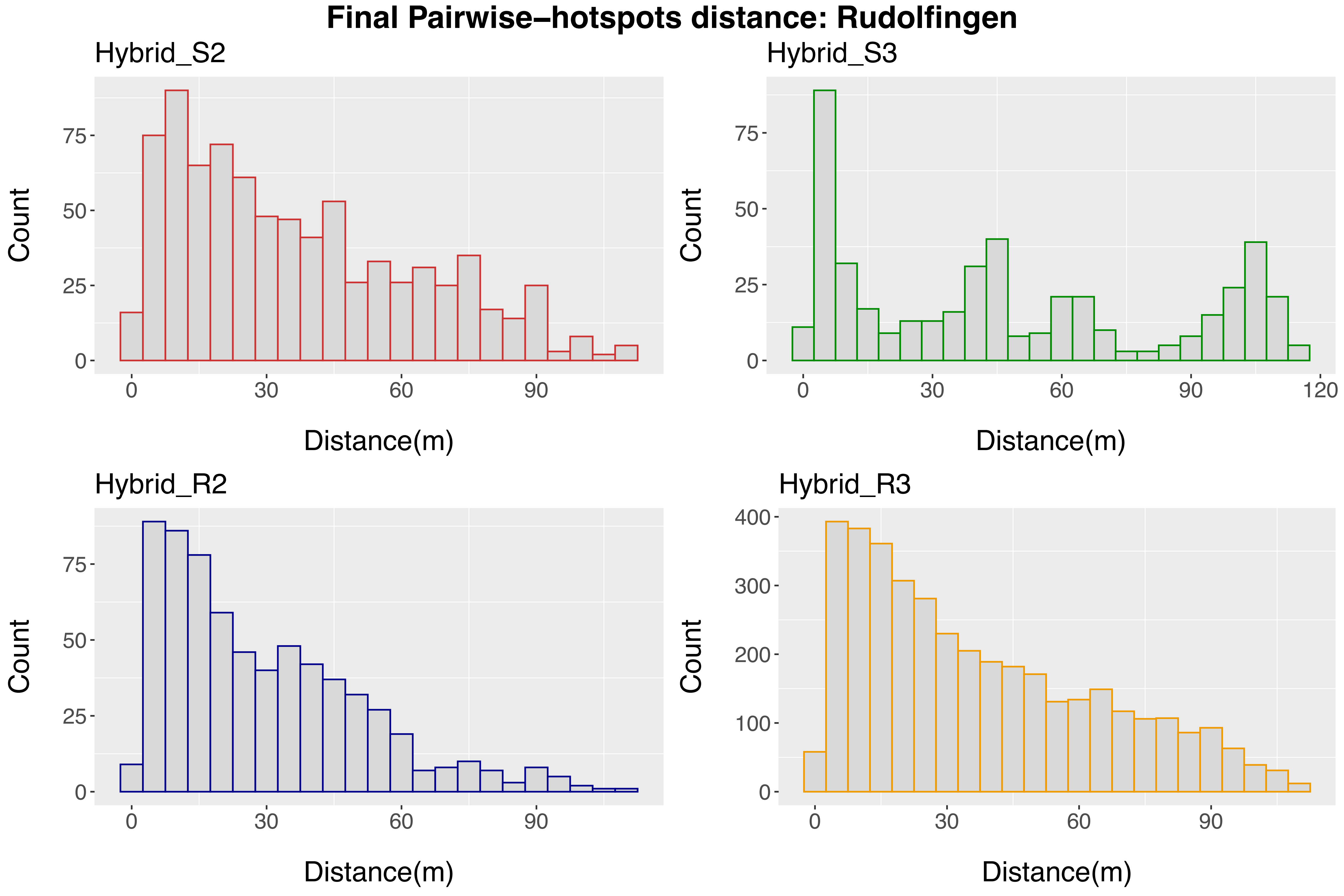
Final Pairwise–hotspots distance: Rudolfingen
Hybrid_S2
Hybrid_S3
Hybrid_R2
Hybrid_R3
Count
Distance(m)
0
25
50
75
100
200
300
400
30
60
90
120

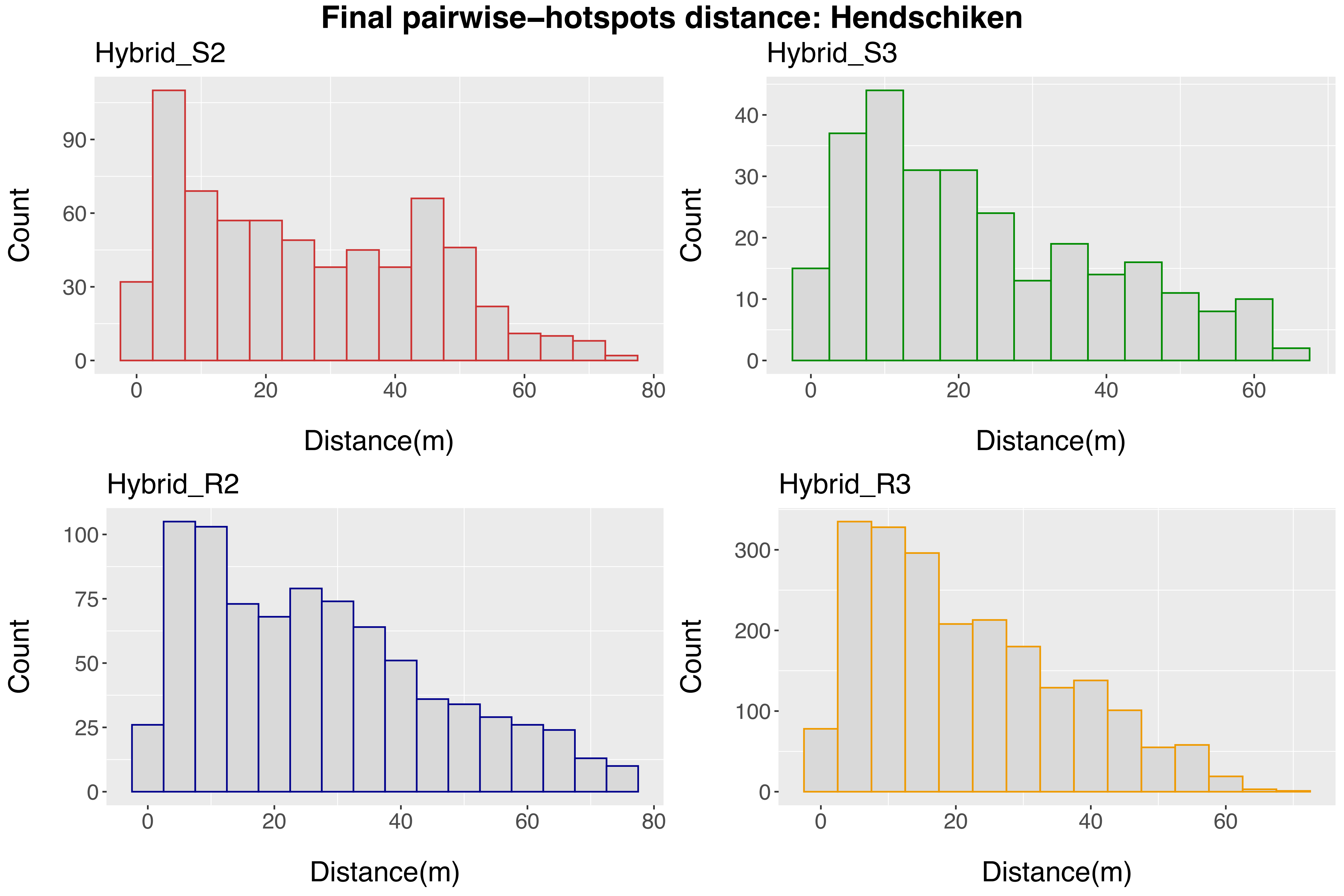
Final pairwise−hotspots distance: Hendschiken
Hybrid_S2
Hybrid_S3
Hybrid_R2
Hybrid_R3
Count
Distance(m)
0
20
40
60
80
30
90
10
100
25
50
75
200
300

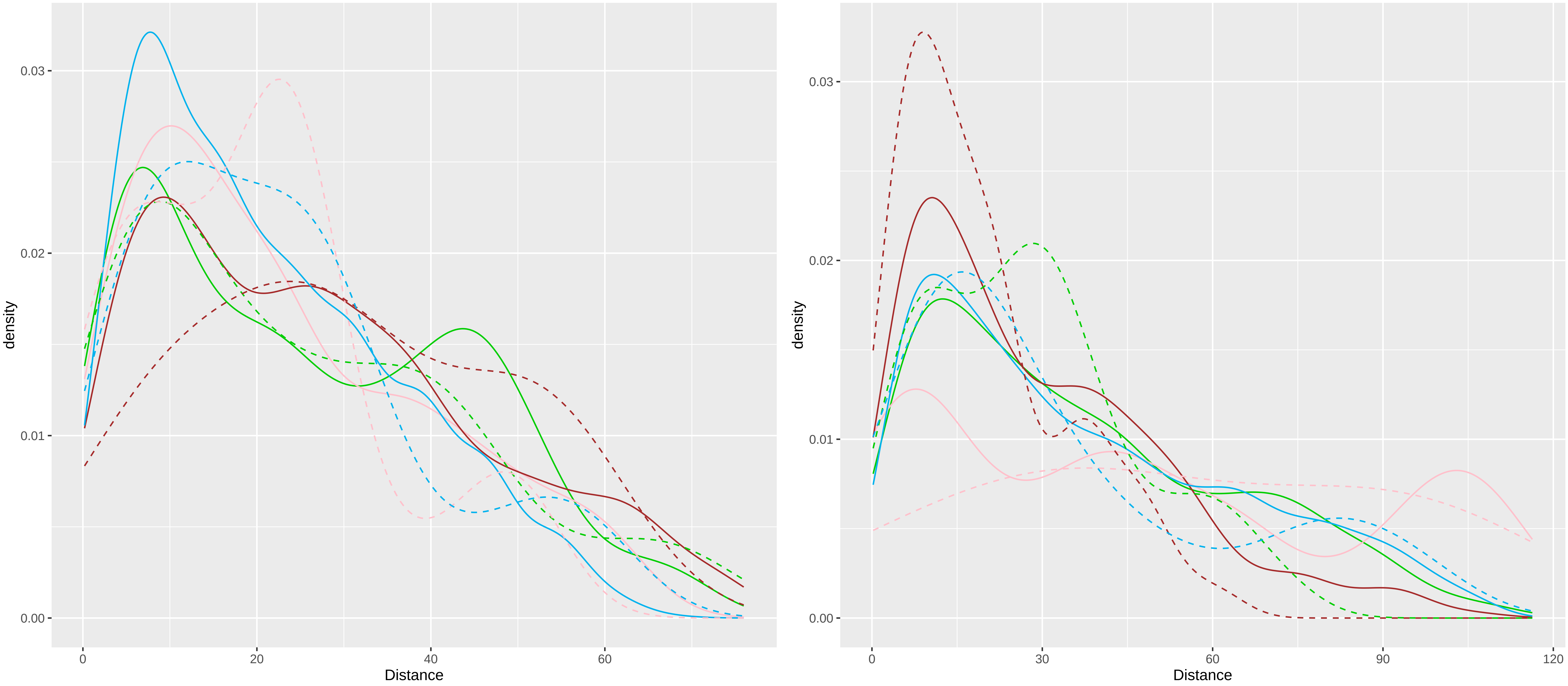

0.03
0.02
0.01
0.00
density
0
20
40
60
Distance
0.03
0.02
0.01
0.00
density
0
30
60
90
120
Distance

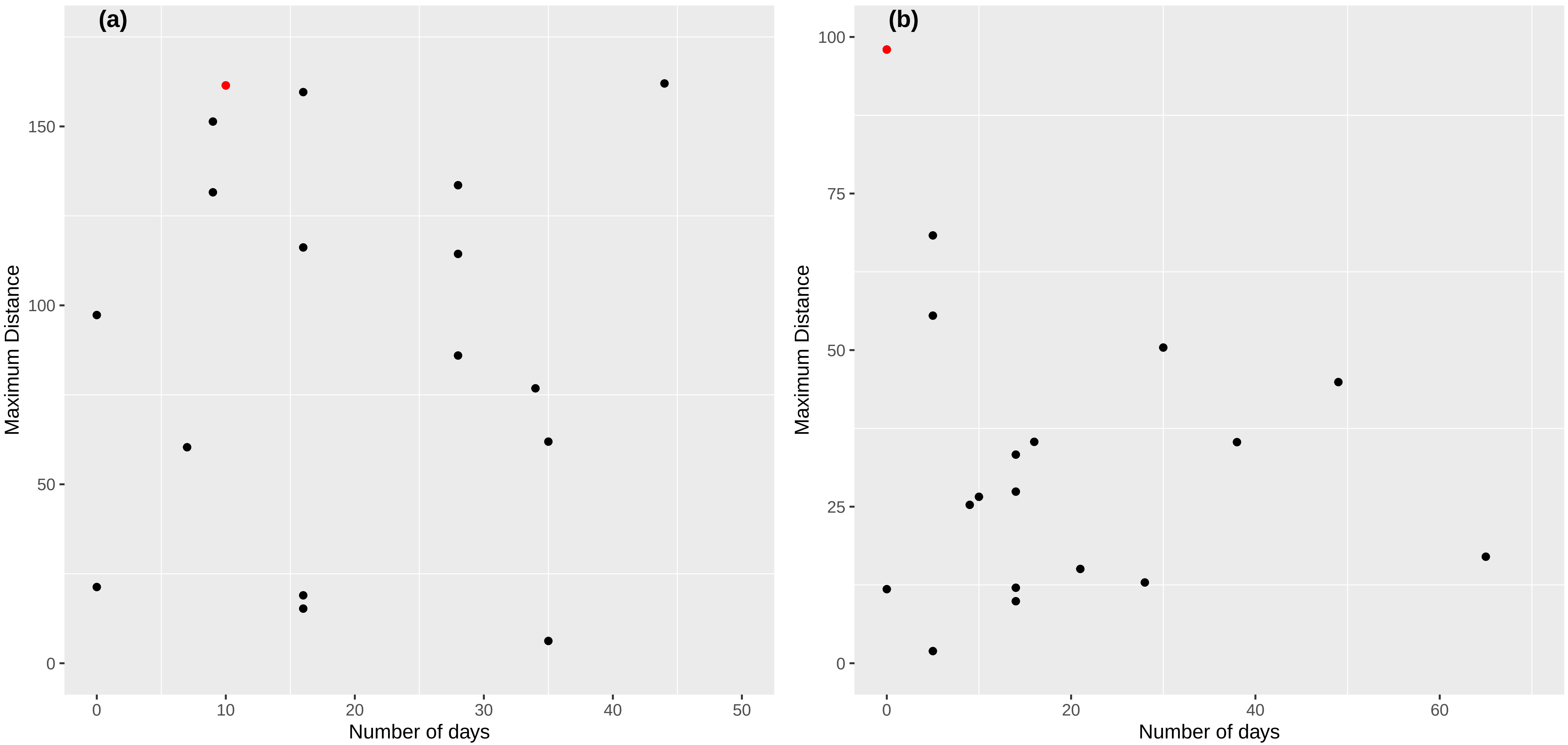

(a)
Maximum Distance
0
50
100
150
0
10
20
30
40
50
Number of days
(b)
Maximum Distance
0
25
50
75
100
0
20
40
60
Number of days
